# Dynamic Changes in the Urinary Proteome of Normal Pregnant Women and Their Correlation with Fetal Developmental Progression -- A Methodological Exploration Based on “One-versus-Many” Urinary Proteomic Comparative Analysis

**DOI:** 10.64898/2026.07.16.738857

**Authors:** Zheng Meilin, Su Yan, Bao Yijin, Sun Wei, Gao Youhe

**Author notes:** Author Biography: Zheng Meilin (2002—), female, Master’s degree candidate, research focus: urinary biomarkers. Corresponding Author: Youhe Gao, Professor, Doctoral Supervisor, Director of the Gene Engineering Drug and Biotechnology Beijing Key Laboratory, College of Life Sciences, Beijing Normal University, Beijing 100875, China.; Wei Sun,Core Facility of Instrument, Institute of Basic Medical Sciences, Chinese Academy of Medical Sciences, School of Basic Medicine, Peking Union Medical College, Beijing, China.

## Abstract

This study employed a “one-versus-many” (a single pregnant woman compared with multiple non-pregnant women) urinary proteomic comparative framework to examine whether fetal development-related signals can be captured through changes in the urinary proteome under conditions of limited sample size. The experimental group consisted of urinary proteomic data from three women with normal pregnancies (R6, R15, R16) at three gestational time points (∼6-8 weeks, 22-24 weeks, and 32-34 weeks; Wang et al., 2022), while the control group consisted of urinary proteomic data from six healthy non-pregnant women (Bao & Gao, ChinaXiv: 202302.00108v2). A total of nine independent “1 vs. 6” differential protein analyses and DAVID GO Biological Process enrichment analyses (P < 0.05) were performed. The results showed that all three pregnant women exhibited a large number of differentially expressed proteins at each time point, and all enriched GO BP terms highly relevant to concurrent fetal organ development (nervous system, lung, eye, ear, kidney, etc.). This suggests that the pregnancy urinary proteome can reflect fetal development signals, corroborating the findings reported by Wang et al. (2025) in a rat model. This study demonstrates that the one-versus-many comparative approach maintains high sensitivity under small-sample conditions and can provide a methodological reference for personalized pregnancy medicine.

## 1. Introduction

Urine is the final product of glomerular filtration and tubular reabsorption of plasma by the kidneys, and contains abundant protein information derived from the kidneys, urinary tract, and various systems of the body^[1][2]^. Compared with blood, urine collection offers the advantages of being non-invasive and repeatable, and because urine need not maintain internal homeostasis, it responds more sensitively to subtle physiological changes in the body^[2]^. Numerous studies in recent years have demonstrated that the urinary proteome can detect differentially expressed proteins of diagnostic or monitoring value in various diseases, including autism, Parkinson’s disease, glioma and hepatocellular carcinoma^[3][4][5][6]^, indicating broad potential for biomarker discovery.

Pregnancy is a special period of systemic physiological remodeling in women, involving the coordinated adaptation of multiple systems including endocrine, immune, and metabolic functions^[9]^. Studies have already investigated the dynamic changes of the urinary proteome during pregnancy: Zheng et al. (2013) systematically compared the urinary proteomes of pregnant and non-pregnant women and revealed marked differences in the pregnancy urinary proteome^[1]^; Wang et al. (2022), in a multi-gestational-stage urine study of 84 pregnant women, found that the urinary proteome not only reflects the physiological changes of normal pregnancy but also enables early prediction of gestational diabetes mellitus (GDM, AUC = 0.91) and spontaneous abortion (SA, AUC = 0.81) at the 6 – 8 weeks gestational time point^[10]^; Tang and Gao (2022) further confirmed in a rat model that the urinary proteome can capture temporal signals of multi-organ development of the fetal heart, lung, liver, and pancreas, with time points highly consistent with embryological observations^[11]^; Wang et al. (2025) refined these findings by collecting daily urine samples from pregnant rats^[12]^.

The conventionally used group-based analysis employs a “many-versus-many” statistical framework for differential protein screening. However, the averaging of cross-individual signals in group-based analysis may mask sensitive signals of important biological significance at the individual level. Existing evidence indicates that the inter-individual biological variation in the urinary proteome of healthy populations is extremely large. Shao, Zhao, and Gao et al. (2019), in a systematic analysis of the urinary proteome of healthy adults, found that the median biological coefficient of variation (CV) of protein abundance between individuals was as high as 0.60 (approximately 60%)^[14]^. Against such a high level of individual variability, group-based differential protein analysis relying on population means inevitably tends to retain common population signals while discarding individual-specific sensitive signals. Moreover, the timing of fetal organ development itself exhibits marked physiological inter-individual variation: Sato and Miyasaka (2019), analyzing longitudinal ultrasound data from 801 normal singleton pregnancies, revealed three distinctly different trajectory patterns of fetal estimated-weight growth velocity, namely decelerating growth (14.7%), stable growth (73.9%), and accelerating growth (11.4%). These trajectory differences could not be readily explained by known factors such as maternal age, height, pre-pregnancy BMI, or fetal sex^[15]^. Even in completely normal pregnancies, the progression of fetal organ development may differ in phase among individuals — at a given gestational week, one pregnant woman may have already entered an active signal peak of a specific organ’s development, whereas in another woman this peak may not yet have appeared or may have already subsided.

Against this background, we adopted the research paradigm of “personal omics profiling” in the field of precision medicine — using the individual as a baseline to longitudinally track the temporal changes of molecular signals^[16]^. The one-versus-many analytical approach has been validated for detecting individual-specific pathways at the single-sample level that are missed by group-based analysis^[17]^. In this study, by replacing the traditional group-based comparison with a one-versus-many comparative strategy, we aimed to preserve individual-specific developmental information and capture sensitive pregnancy signals that might be diluted by group-based analysis. The experimental group consisted of urinary proteomic data from three healthy pregnant women (R6, R15, R16, derived from the normal-control group in Wang et al., 2022) at three time points, and the control group consisted of urine data from six healthy non-pregnant women in Gao’s research group^[13]^. We constructed a one-versus-many differential protein analysis system and performed GO Biological Process (BP) enrichment analysis on the differential proteins of each pregnancy stage using the DAVID database.

## 2. Materials and Methods

### 2.1. Sample Sources

The experimental-group urine data were derived from the pregnancy urinary proteomic study published by Wang et al. (2022)[10], which was approved by the Ethics Committee of Peking Union Medical College Hospital (No. ZS-976) and conducted in accordance with the Declaration of Helsinki. We used data from pregnant women R6, R15, and R16 in the normal-pregnancy control group of that study; none of the three women developed GDM or experienced spontaneous abortion during the entire follow-up period, representing a normal physiological pregnancy process. Urine samples were collected three times: the first collection at a time point during 6–8 weeks of gestation, the second at a time point during 22–24 weeks, and the third at a time point during 32–34 weeks, using first-morning midstream urine, for a total of 9 samples.

The control-group data were derived from the study published by Bao et al. of our research group^[13]^. Baseline urine samples (T0 time point) collected before massage from six healthy non-pregnant female graduate students were used as the non-pregnant control group. All six subjects were 22 – 26 years old, with no pregnancy history, no chronic disease, and no recent medication history.

### 2.2. Urinary Protein Extraction and Enzymatic Digestion

Protein extraction and enzymatic digestion for the experimental group followed the method of Wang et al. (2022)^[10]^: 10 mL of first-morning midstream urine was taken, proteins were extracted by acetone precipitation after centrifugation, and digestion was performed using the filter-aided sample preparation (FASP) method; the reducing reagent was DTT, the alkylating reagent was IAA, and digestion was carried out with trypsin (Trypsin Gold, Promega) at 37°C for 8 h, followed by desalting on an HLB column (Waters) and storage at −80°C until use.

The control-group sample processing procedure was as follows: 4 mL of first-morning midstream urine was centrifuged at 4°C and 12,000 × g for 30 min, and the supernatant was collected; after a second centrifugation at 12,000 × g for 30 min, the supernatant was taken and lysis buffer was added (8 mol/L urea, 50 mmol/L Tris-HCl, pH 8.5, containing 2 mol/L thiourea, 25 mmol/L NH4HCO3, and 20 mmol/L DTT), and protein concentration was determined by the Bradford method. Digestion was performed by the FASP method (10 kDa cut-off, PALL membrane): DTT (25 mmol/L) was added to UA (8 mol/L urea, 0.1 mol/L Tris-HCl, pH 8.5) and reduced at 37°C for 1 h, followed by alkylation with IAA (50 mmol/L) in the dark for 30 min; after buffer exchange with 25 mmol/L NH4HCO3, Trypsin Gold (50:1 w/w) was added and digestion was performed overnight at 37°C; the sample was desalted on an HLB column (Waters) and stored frozen at −80°C^[13]^.

### 2.3. Liquid Chromatography–Tandem Mass Spectrometry (LC-MS/MS) Analysis

Mass spectrometry analysis of the experimental group was performed on an Orbitrap Fusion Lumos Tribrid mass spectrometer (Thermo Fisher Scientific, USA) coupled with an EASY-nLC 1000 (Thermo Fisher Scientific). Peptides were fractionated by high-pH reversed-phase liquid chromatography (RPLC) and acquired in DDA mode to build a spectral library; targeted DIA acquisition used a scan range of m/z 400–900, with resolving power of 120,000 (MS1) and 30,000 (MS2), and an HCD fragmentation energy of 32%. Detailed instrument parameters are described in Wang et al. (2022)^[10]^.

Mass spectrometry analysis of the control group was performed on the same model of Orbitrap Fusion Lumos Tribrid mass spectrometer coupled with an EASY-nLC 1200 (Thermo Fisher Scientific). Peptides were first acquired three times in DIA mode, with simultaneous DDA acquisition for library construction. The DIA method was set with reference to BCA quantification results, at a sample concentration of 0.5 μg/μL, an injection volume of 9 μL per run, and a total injection amount of approximately 500 ng; a 90-min gradient elution was used (mobile phase A: 0.1% formic acid in water; mobile phase B: 80% acetonitrile containing 0.1% formic acid); iRT standards (Biognosys) were added for retention-time correction. Detailed parameters are described in Bao et al.^[13]^.

### 2.4. Protein Identification and Quantification

Raw data analysis of the experimental group: DDA data were searched using Proteome Discoverer (PD) 2.1 against the UniProt Human database (version 2017_09), with peptide FDR < 1%, protein FDR < 1%, and a maximum of 2 missed cleavages allowed; DIA data were then imported into Spectronaut Pulsar 14.10, and a spectral library was built based on the above DDA search results for DIA quantitative analysis, requiring the identification of at least 2 unique peptides per protein, with a q-value < 0.01 (FDR 1%). See Wang et al. (2022)^[10]^ for details.

Raw data analysis of the control group: DDA data were first searched using Proteome Discoverer 2.1 (PD 2.1) against the UniProt Human database, and the search results were used to build the spectral library required for DIA analysis. DIA quantitative analysis was also performed using Spectronaut X, with a q-value < 0.01, i.e., FDR 1%^[13]^.

### 2.5. One-versus-Many Comparative Analysis of Differential Proteins

Data integration and analysis in this study: the experimental-group subjects R6/R15/R16 were each sampled at a time point within 6–8 weeks, 22–24 weeks, and 32–34 weeks, with three technical replicates per sample. The first sampling is denoted R6-T1, R15-T1, R16-T1; the second sampling R6-T2, R15-T2, R16-T2; and the third sampling R6-T3, R15-T3, R16-T3. The control group consisted of six non-pregnant women (C1–C6), with three technical replicates per sample.

The raw mass spectrometry data of the experimental and control groups were searched and quantified in four batches using Spectronaut X software. The search reference databases were iRT peptides-fusion and UniProt Human, with q < 0.01, i.e., FDR 1%.

The batches were set up as follows: Batch 1 (R6/R15/R16 each at the 6–8 weeks time point + 6 controls), Batch 2 (R6 at the 22–24 weeks and 32–34 weeks time points + 6 controls), Batch 3 (R15/R16 each at the 22–24 weeks time point + 6 controls), and Batch 4 (R15/R16 each at the 32–34 weeks time point + 6 controls). All four batches used identical Spectronaut X search parameters, and each batch contained the same six non-pregnant control samples as an inter-batch reference, so that batch effects could be avoided in subsequent differential analysis through within-batch paired comparisons.

For each sampling, the protein quantification data of the experimental group were independently compared with those of the six non-pregnant women in the control group (C1–C6), constructing a “1 vs. 6” single-sample-versus-multiple-sample differential analysis, for a total of nine independent comparisons. The differential protein screening procedure was as follows:

1. Calculation of the control-group mean: each of the six control subjects had three technical replicates measured (18 data points in total); the mean of the three replicates for each subject was first calculated, and then the mean of the six subjects’ means was calculated and recorded as the “control-group mean” for each protein.
2. Calculation of the experimental-group mean: each pregnant woman’s sample was measured in three technical replicates, and the average of the three quantitative values was taken as the “experimental-group mean” for each protein.
3. Fold change (FC) calculation: FC was calculated using the formula FC = experimental-group mean ÷ control-group mean. An FC ≥ 1.5 was defined as up-regulation and FC ≤ 0.67 (i.e., 1/1.5) as down-regulation. FC = 1.5 is a conventionally used threshold in proteomic differential-expression analysis, which balances a certain biological effect size (50% change) with sensitivity. In addition, proteins with a control-group mean of zero or infinity (i.e., detectable only in pregnant women) were also considered to meet the FC screening criterion.
4. Statistical test: Welch’s two-tailed t-test with unequal variance was used to compare the significance of differences between the 18 data points of the control group and the 3 data points of the experimental group. This was implemented using the TTEST function in Microsoft Excel (formula: TTEST(control-group data range, experimental-group data range, 2, 3), where parameter 2 denotes a two-tailed test and parameter 3 denotes the assumption of unequal variance).
5. Differential protein screening criteria: a protein was selected if it simultaneously satisfied P < 0.05 and (FC ≥ 1.5 or FC ≤ 0.67), or if it was undetectable in the control group (mean zero/infinity) but detectable in pregnant women.

The following methodological issues need to be clarified:

1. Issue of biological replicates in the experimental group: no biological replicates were set for the pregnant-woman samples in the experimental group; the data used three technical replicates. Under the Welch t-test framework, the control-group variance originates from the biological variation of six independent individuals, whereas the experimental-group variance derives only from technical replicate error (usually far smaller than biological variation). Therefore, the statistical power of this study may be either conservative or liberal, depending on the magnitude and direction of the control-group variance. This is also one of the methodological limitations of the one-versus-many analytical approach.
2. Note on multiple hypothesis testing correction: in the differential protein screening step (nine independent comparisons, each involving t-tests of thousands of proteins), we simultaneously applied the Benjamini–Hochberg (BH) FDR correction (step-up method, 1995) to the raw P-values for multiple hypothesis testing correction. The correction results showed that among the nine comparisons, a total of 22,679 proteins had a raw P < 0.05, and 22,469 high-confidence differential proteins remained after BH correction at FDR < 0.05, an overall retention rate of 99.1%. The post-correction retention rates for each comparison were as follows: R6-T1 (2,279/2,279, 100%), R6-T2 (2,677/2,677, 100%), R6-T3 (2,897/2,969, 97.6%), R15-T1 (1,305/1,305, 100%), R15-T2 (2,849/2,849, 100%), R15-T3 (2,892/2,892, 100%), R16-T1 (1,760/1,760, 100%), R16-T2 (2,942/2,942, 100%), and R16-T3 (2,868/3,006, 95.4%). Among the nine comparisons, only R6-T3 and R16-T3 showed a small number of proteins removed by FDR correction (decreased by 2.4% and 4.6%, respectively), while all proteins in the other seven comparisons passed FDR correction. The fundamental reason why FDR correction had minimal impact on the results is that the three technical replicates of the experimental group had identical quantitative values (3 technical replicates), resulting in zero within-group variance and extremely small Welch t-test P-values (mostly on the order of 10⁻⁶ –10⁻¹⁰), which were essentially unaffected after BH correction. Based on the above FDR-corrected results, this study defined the final differential proteins as those with FDR < 0.05 and meeting the FC screening criteria (FC ≥ 1.5 or FC ≤ 0.67).

### 2.6. Bioinformatics Analysis

The UniProt ID lists of differential proteins from each sample were submitted to the DAVID database (version 6.8, https://david.ncifcrf.gov/) for GO Biological Process (BP) enrichment analysis. The input identifier type was UNIPROT_ACCESSION, the gene list type was Gene List, and GOTERM_BP_DIRECT under the Gene_Ontology category was selected as the enrichment term, with results viewed in Chart mode (displayed as a classic table after functional annotation clustering). A Benjamini–Hochberg-corrected P < 0.05 was used as the threshold for significant enrichment. The background reference gene set was the Homo sapiens whole genome (DAVID default setting).

## 3. Results and Analysis

The experimental-group data in this study were derived from the pregnancy urinary proteome study of Wang et al. (2022), and the control-group data were derived from the baseline urinary proteome data of six healthy non-pregnant women established by Bao et al. The three women with normal pregnancies, R6, R15, and R16, were each sampled three times: the first sampling at a time point during 6–8 weeks of gestation (hereafter denoted R6-T1, R15-T1, R16-T1), the second at a time point during 22–24 weeks (hereafter R6-T2, R15-T2, R16-T2), and the third at a time point during 32–34 weeks (hereafter R6-T3, R15-T3, R16-T3). Using a one-versus-many comparative analysis framework, DAVID GO BP enrichment analysis (P < 0.05) was performed on the differential proteins of the above nine samples. The most important finding of this study corroborates the rat pregnancy urinary proteome study of Wang et al. (2025)—where biological-process signals related to fetal development could be detected in the urinary proteome of rats, the same phenomenon was observed in human pregnancy: each pregnant woman exhibited GO BP terms related to fetal organ development in her urinary proteome at every sampling time point. Across all nine samples, a large number of organ/tissue development or morphogenesis-related terms were identified, covering multiple organ systems including nervous, vascular, cardiac, skeletal, renal, pulmonary, digestive, sensory, and skin.

Most importantly, we clearly identified terms in the category of organ development or morphogenesis in six samples (Table 1), where “No.” denotes the ascending order number of each term within its respective GO BP table, sorted by p-value, corresponding to the ordering in Tables 2 – 10. Many of the biological processes involved in these terms — including placenta blood vessel development, embryonic limb morphogenesis, in utero embryonic development, and regulation of embryonic development — are all related to fetal development and regulation; development of the lung, eye, ear, and kidney, among others, belongs to organ systems that have already completed development in adults, and should not involve continuously active organ morphogenesis in pregnant women, strongly suggesting that the urinary proteome can capture fetal development signals.

**Table 1.**
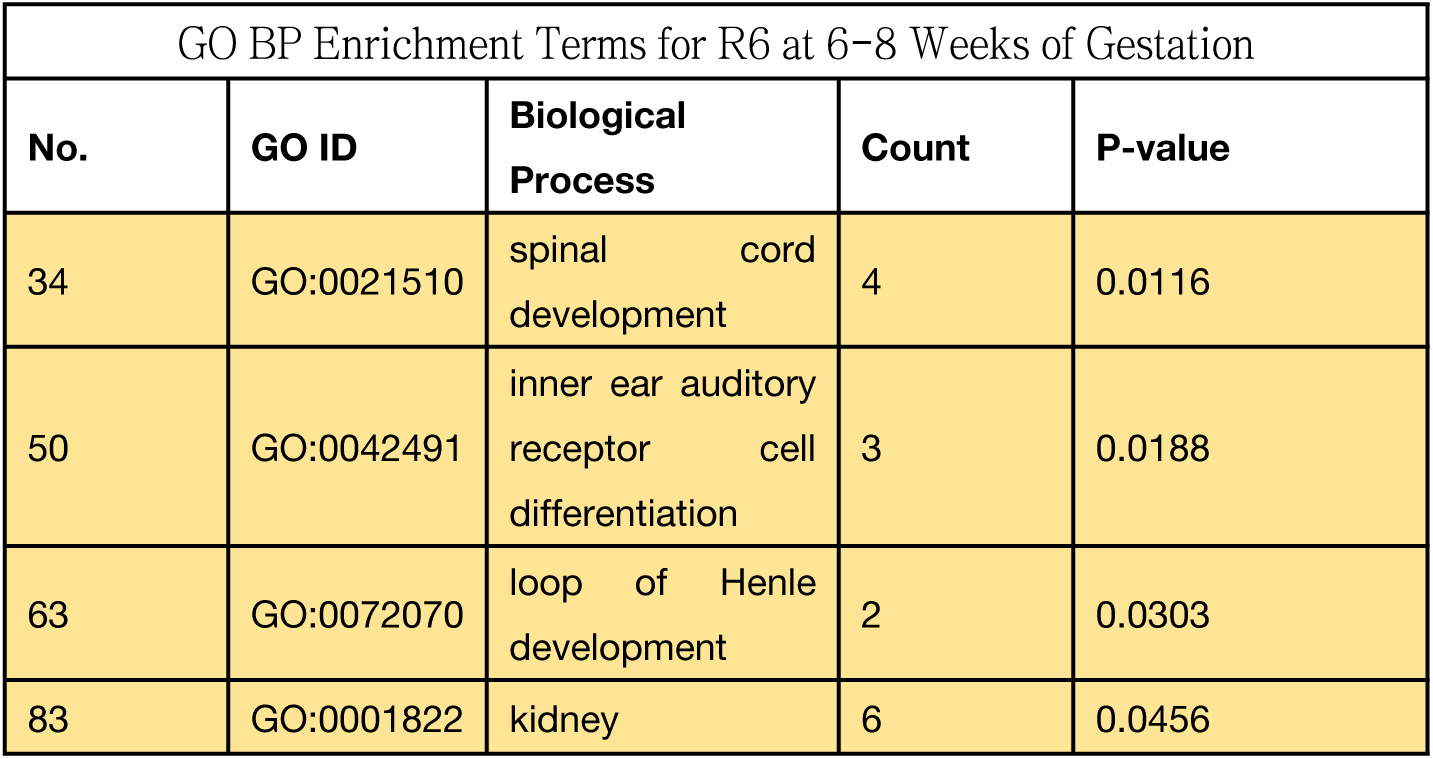
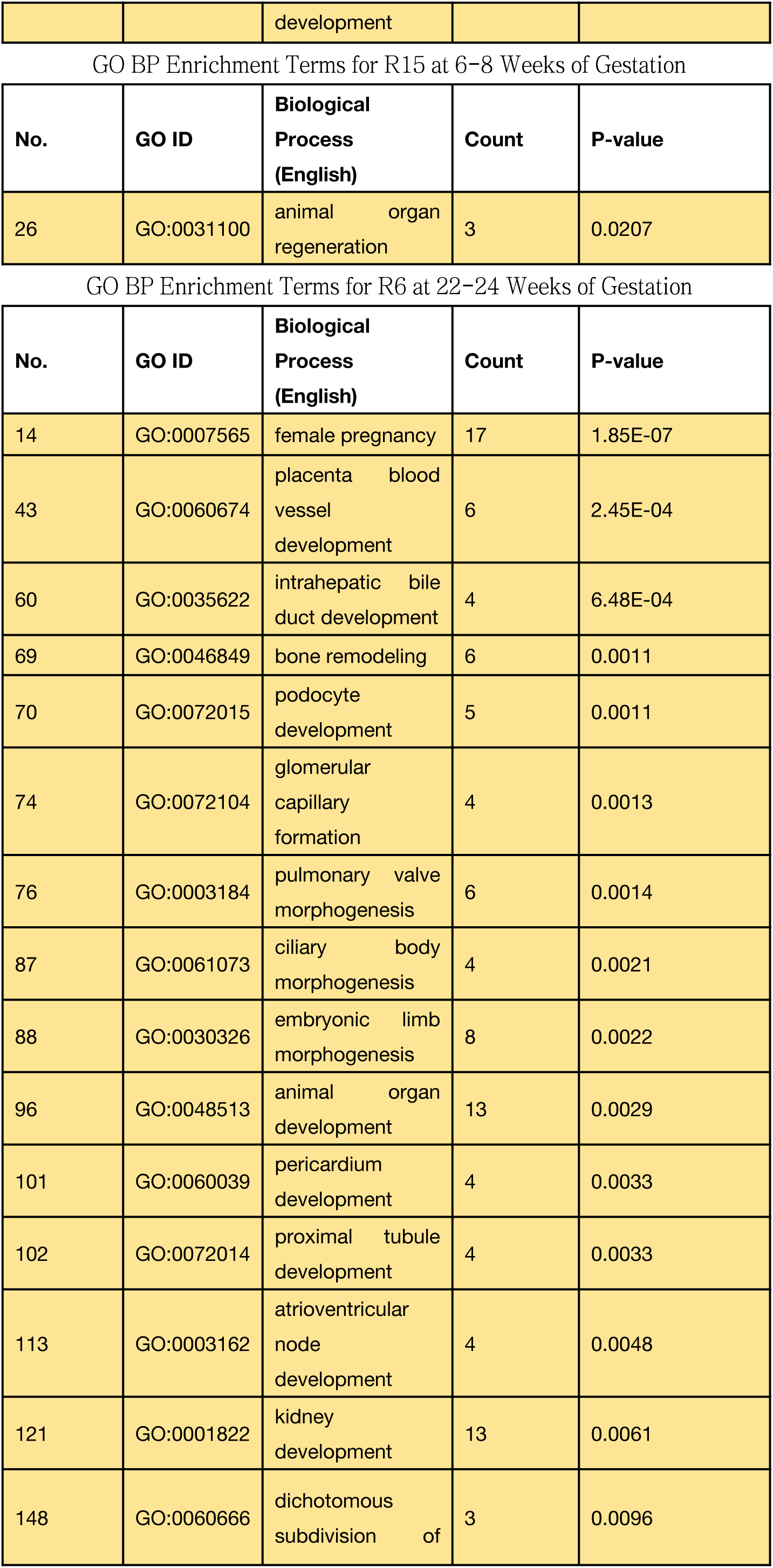
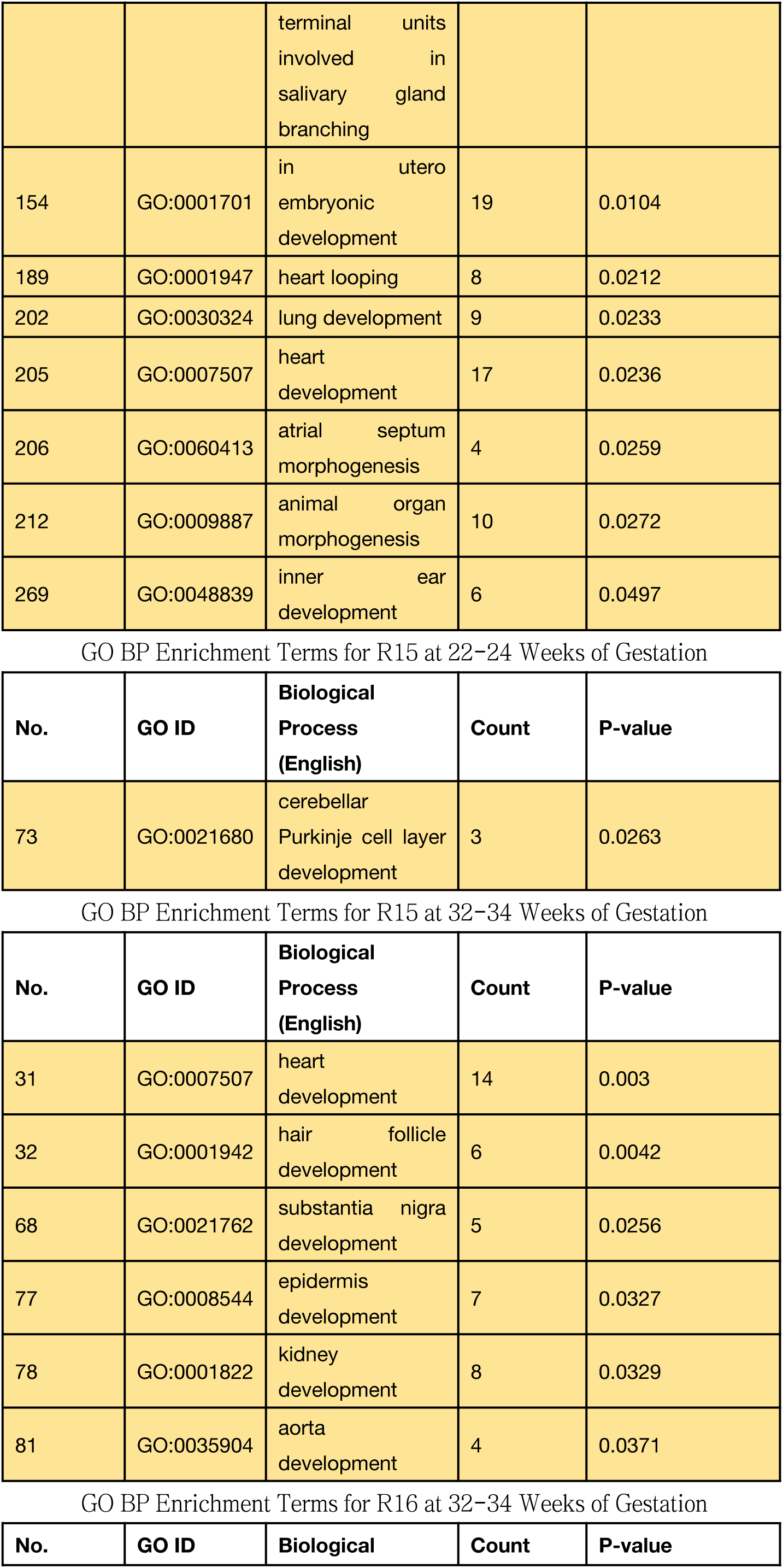
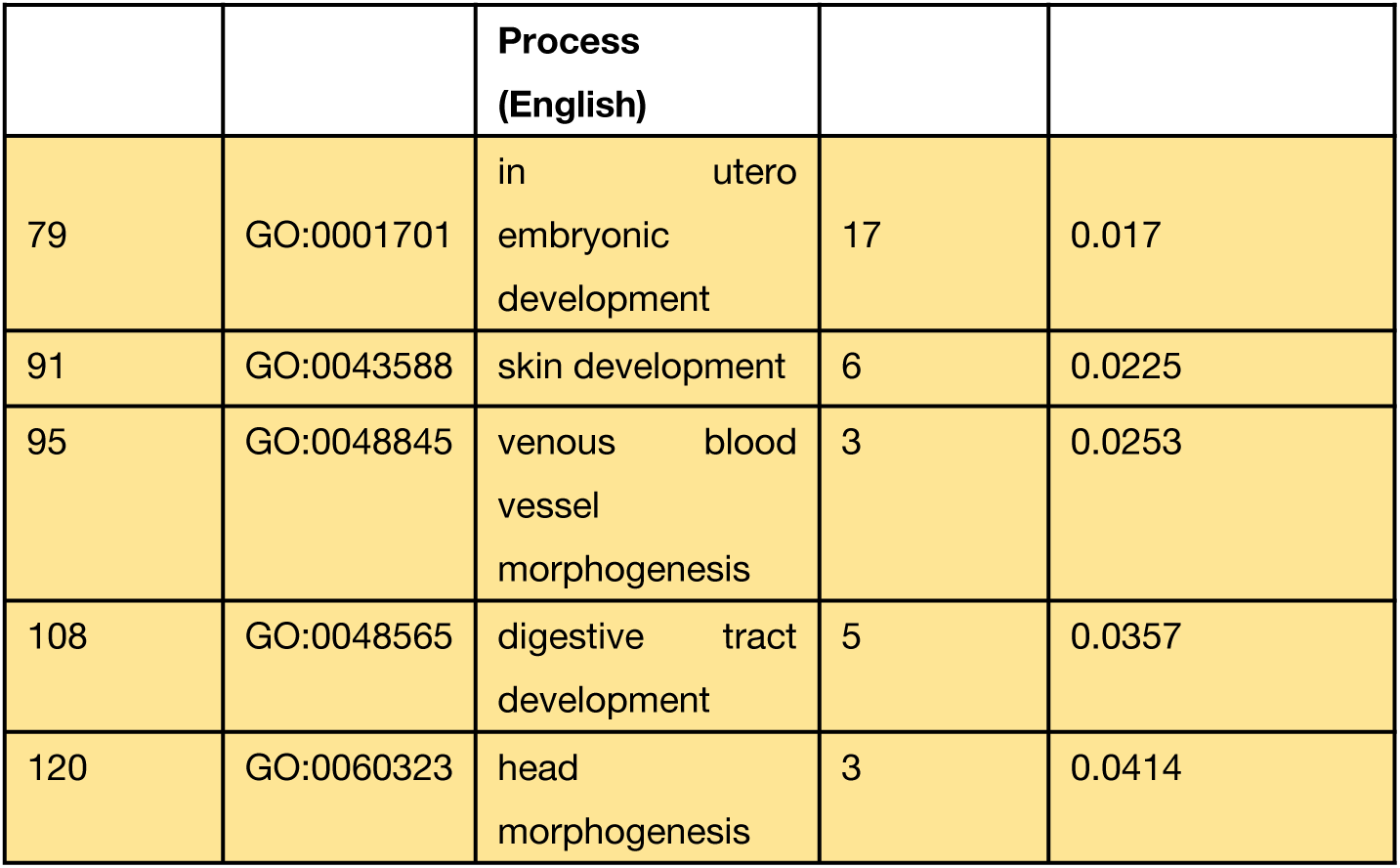
Terms related to organ development or morphogenesis among all GO BP terms.

The main findings at each sampling time point are first summarized by organ system, and then the detailed results for each sample are reported in turn.

Terms related to the cardiovascular system and vascular development were detected at multiple sampling time points in several subjects. Angiogenesis terms appeared at seven time points: R6-T1, R6-T2, R6-T3, R15-T2, R15-T3, R16-T2, and R16-T3; placenta blood vessel development was seen at R6-T2; venous blood vessel morphogenesis at R16-T3; and endothelial cell differentiation and erythrocyte differentiation at R15-T3. Terms related to heart development were seen at R6-T2 and R15-T3: including heart development, pulmonary valve morphogenesis, pericardium development, and atrioventricular node development (all at R6-T2), as well as cardiac muscle cell proliferation (at R16-T2).

Terms related to skeletal and limb development appeared at multiple time points: ossification was seen at R15-T1 and R6-T2; embryonic limb morphogenesis at R6-T2; osteoblast differentiation at R15-T3; and osteoclast differentiation at R6-T3. Terms related to overall embryonic developmental regulation included: in utero embryonic development (seen at R6-T2 and R16-T3), regulation of embryonic development (seen at R15-T2), establishment of left/right asymmetry in lateral mesoderm (seen at R15-T2), and endodermal cell differentiation (seen at R15-T3).

Terms related to urinary system development included kidney development (seen at R6-T1, R6-T2) and podocyte development (seen at R6-T2); terms related to respiratory and digestive system development included lung development (seen at R6-T2), intrahepatic bile duct development (seen at R6-T2), and digestive tract development (seen at R16-T3); terms related to sensory development included inner ear auditory receptor cell differentiation (seen at R6-T1) and inner ear development (seen at R6-T2); and terms related to skin and ectoderm-derived structure development included keratinocyte differentiation (seen at R16-T1), hair follicle development (seen at R15-T3), epidermis development (seen at R15-T3), and epithelial cell differentiation (seen at R15-T3, R16-T3).

Terms related to nervous system development appeared most frequently across all nine sampling time points and constituted the largest and strongest-signal category. R6-T1 detected spinal cord development, neuron migration, neuron projection development, synapse assembly, and central nervous system development; neither R15-T1 nor R16-T1 showed significant neurodevelopmental terms; R6-T2 detected axon guidance and nervous system development; R15-T2 detected cerebellar Purkinje cell layer development; R16-T2 detected neuron migration and axon guidance; R6-T3 detected only a very small number of terms, with neurodevelopmental signals essentially subsided; R15-T3 detected axon guidance, nervous system development, neuron projection development, synapse assembly, retinal ganglion cell axon guidance, and substantia nigra development; and R16-T3 detected commissural neuron axon guidance, nervous system development, axon guidance, and neuron projection development. The above results indicate that the intensity and specific terms of neurodevelopmental signals differed at each sampling time point, and each represents only an independent observation of that time point’s pregnancy urinary proteome.

Most of the biological processes involved in the above terms could hardly originate from the pregnant woman’s own fully developed and mature organs—the placenta in placenta blood vessel development is a fetus-derived organ; embryonic limb morphogenesis, in utero embryonic development, and regulation of embryonic development are all embryo/fetus-specific developmental processes; and the lung, heart, kidney, eye, and ear all belong to organ systems that have completed development in adults, so there should be no continuously active organ morphogenesis in pregnant women. The widespread detection of the above terms across all nine sampling time points strongly suggests that the signals originate from the developing fetus and that the urinary proteome can capture fetal development signals. The observations at each sampling time point are independent and represent only the actual state of the pregnancy urinary proteome at that time point.

The GO BP enrichment results for each subject at each sampling time point are reported below in turn; the complete term data are presented in Tables 2–10, with organ/tissue development terms highlighted.

**Table 2.**
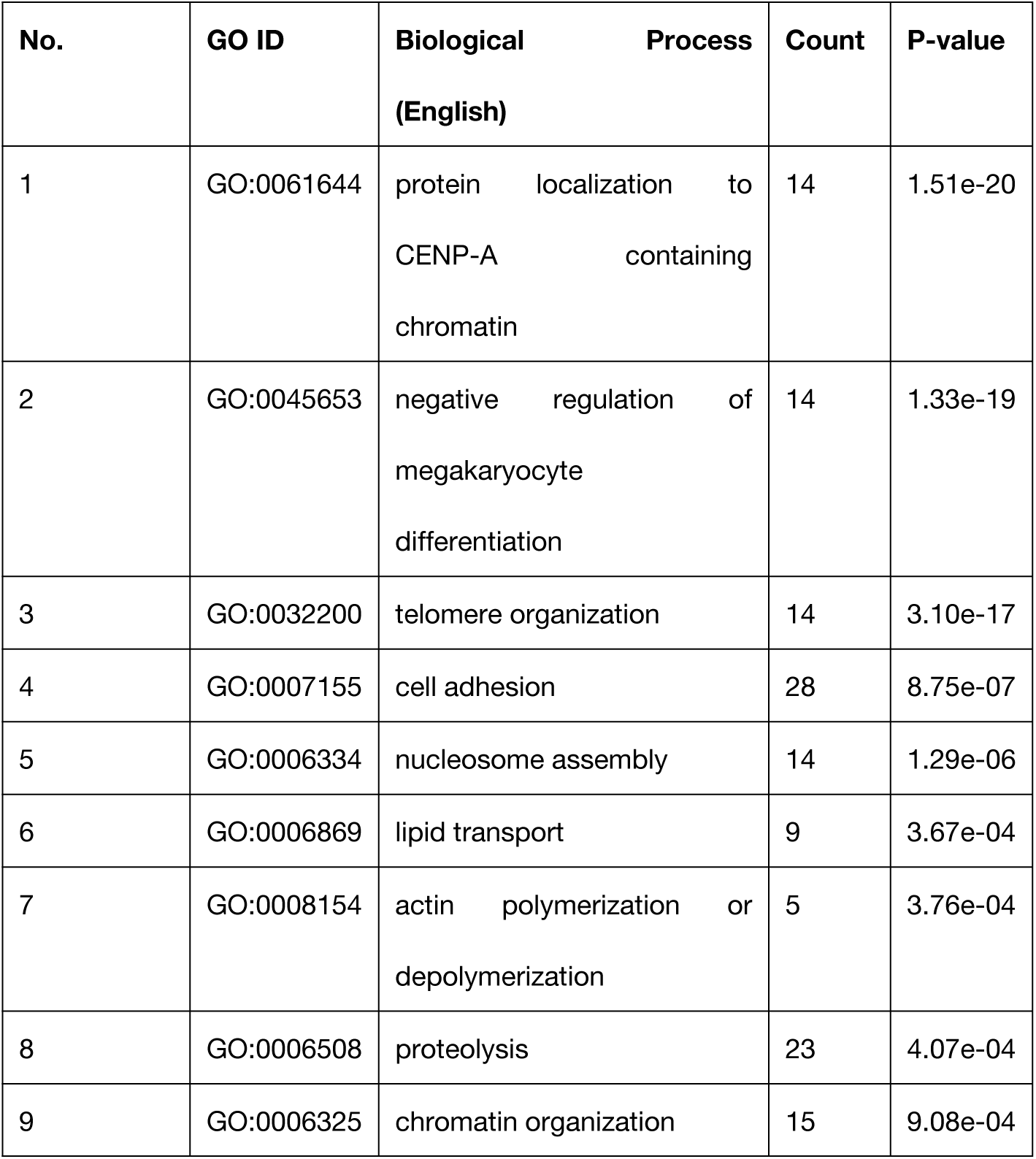
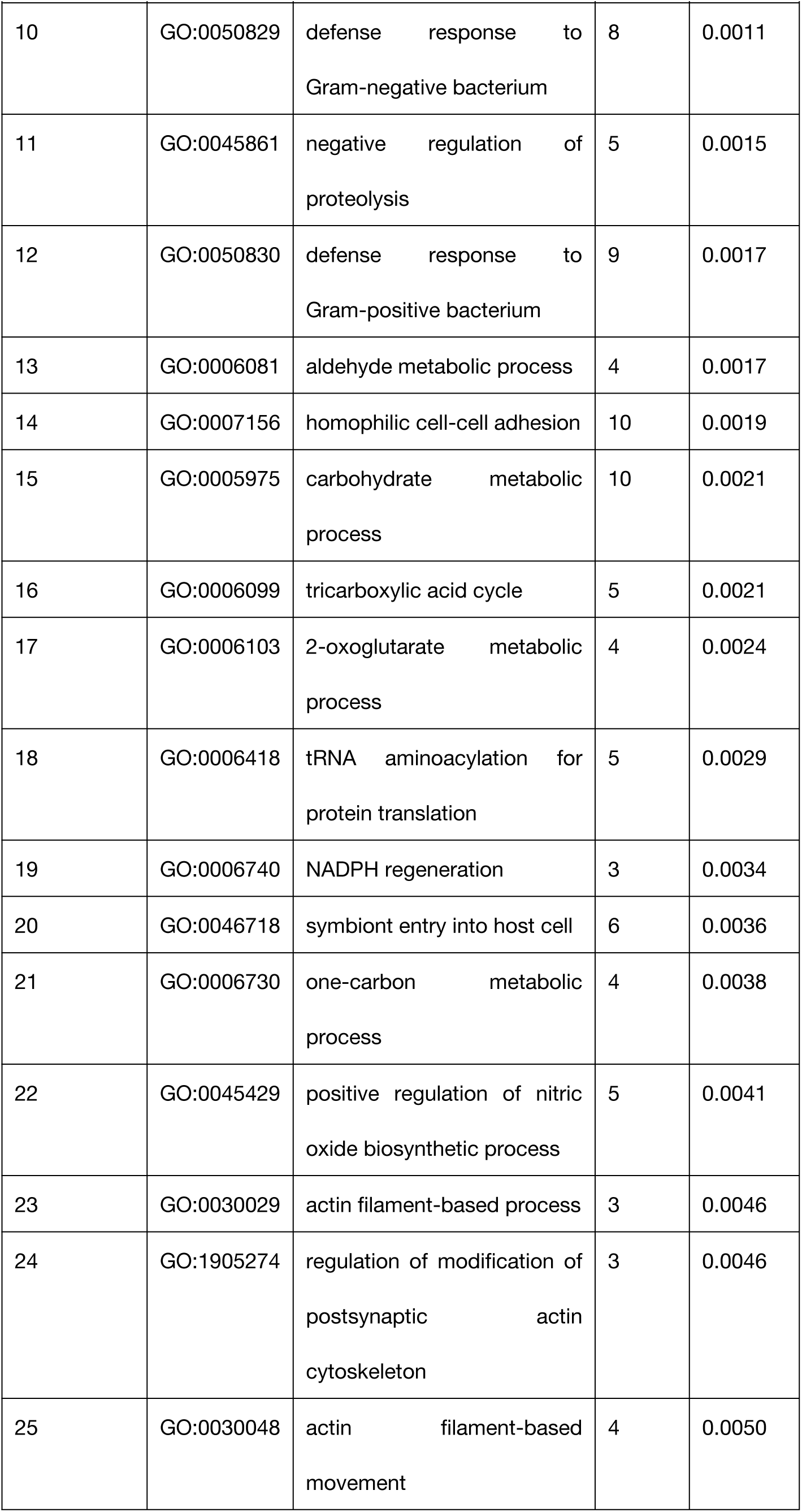
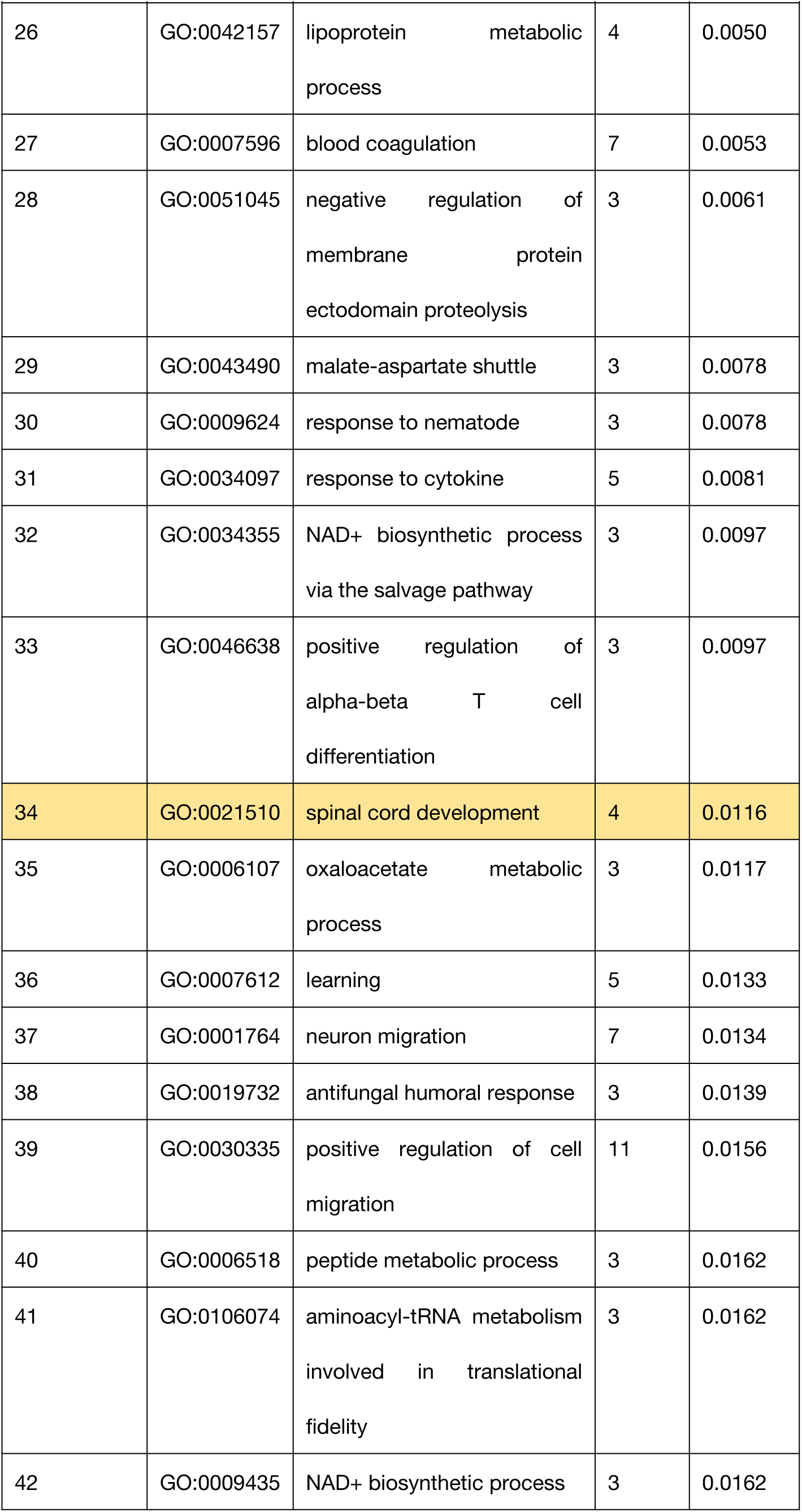
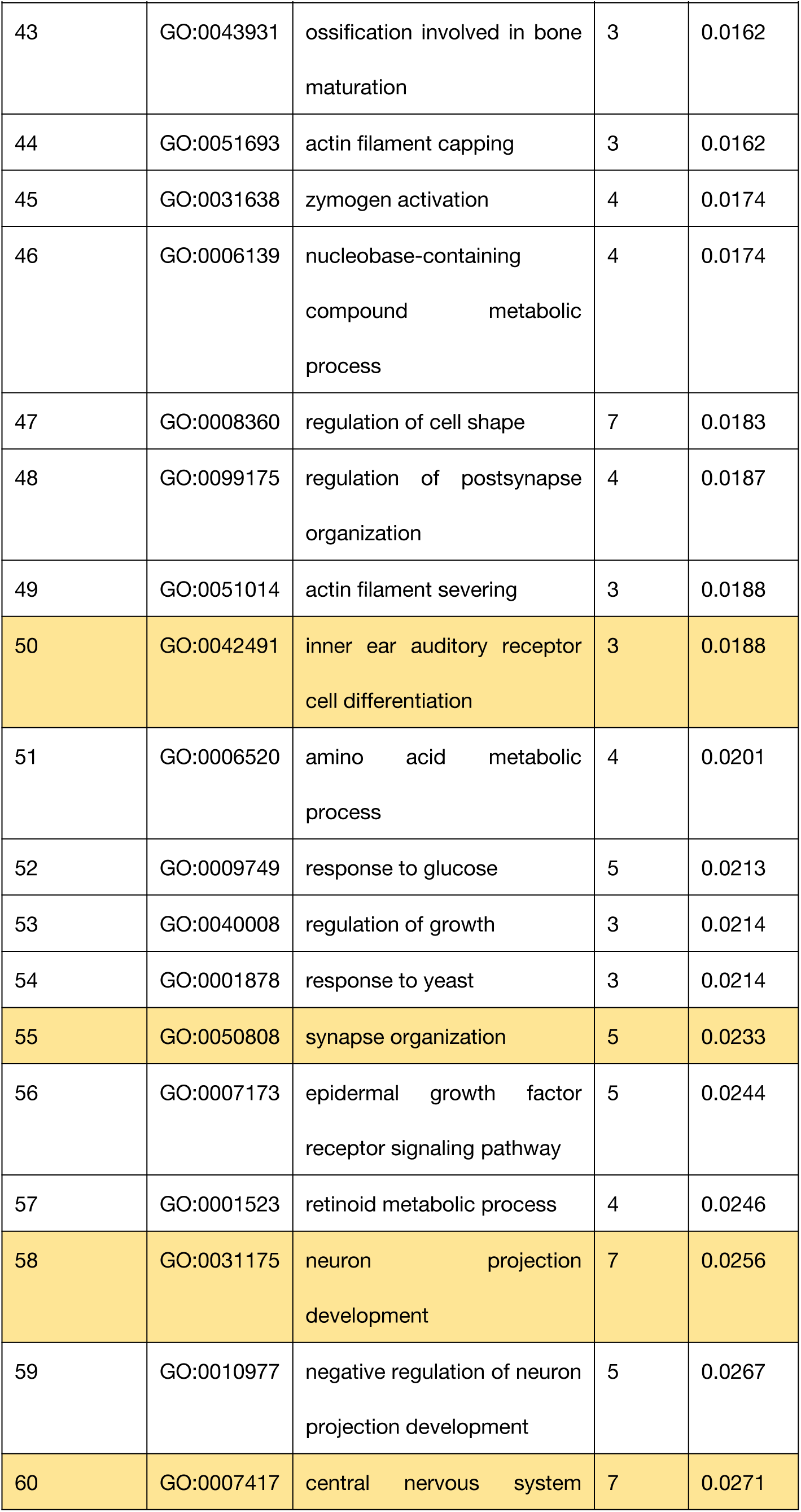
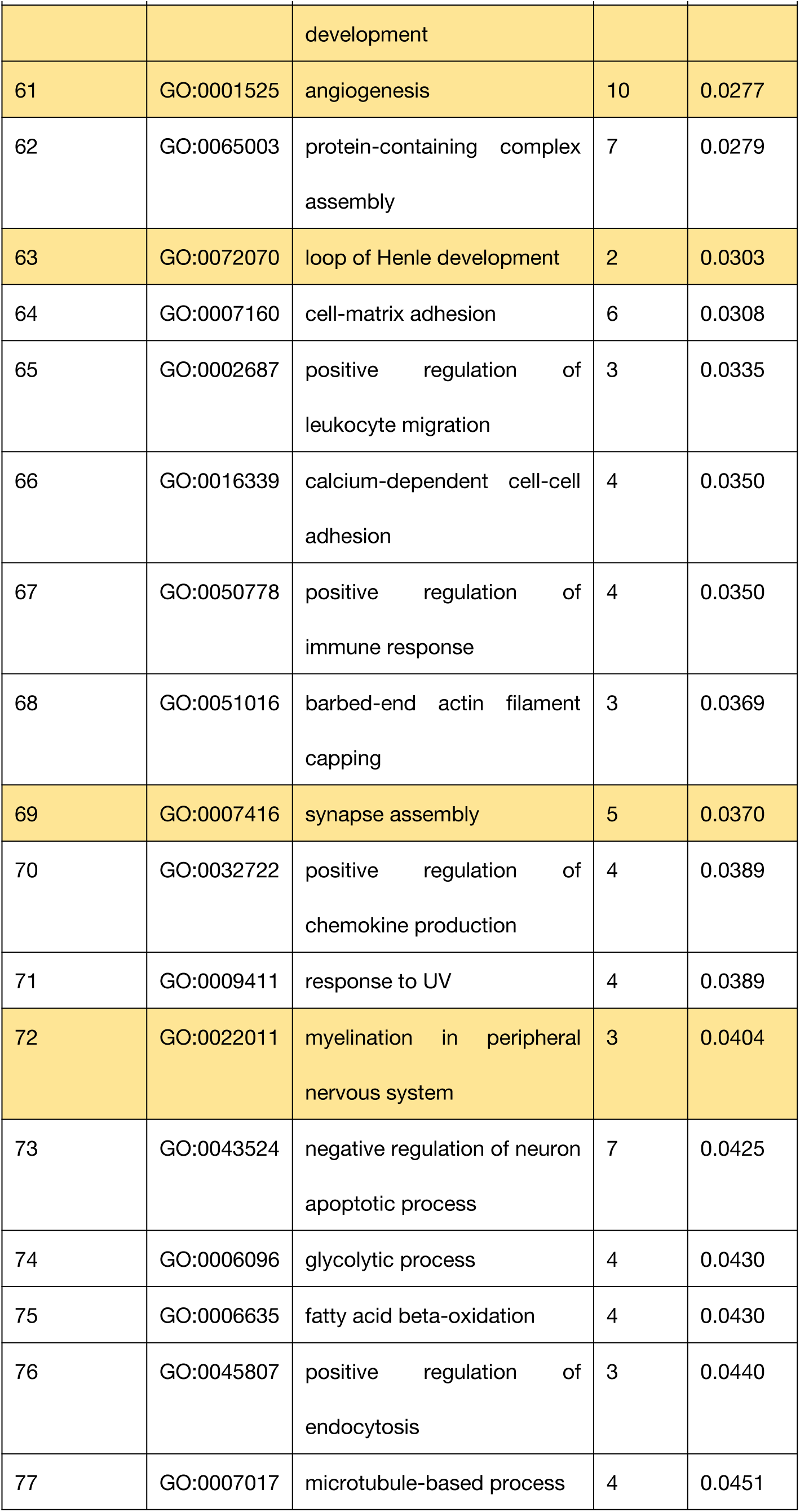
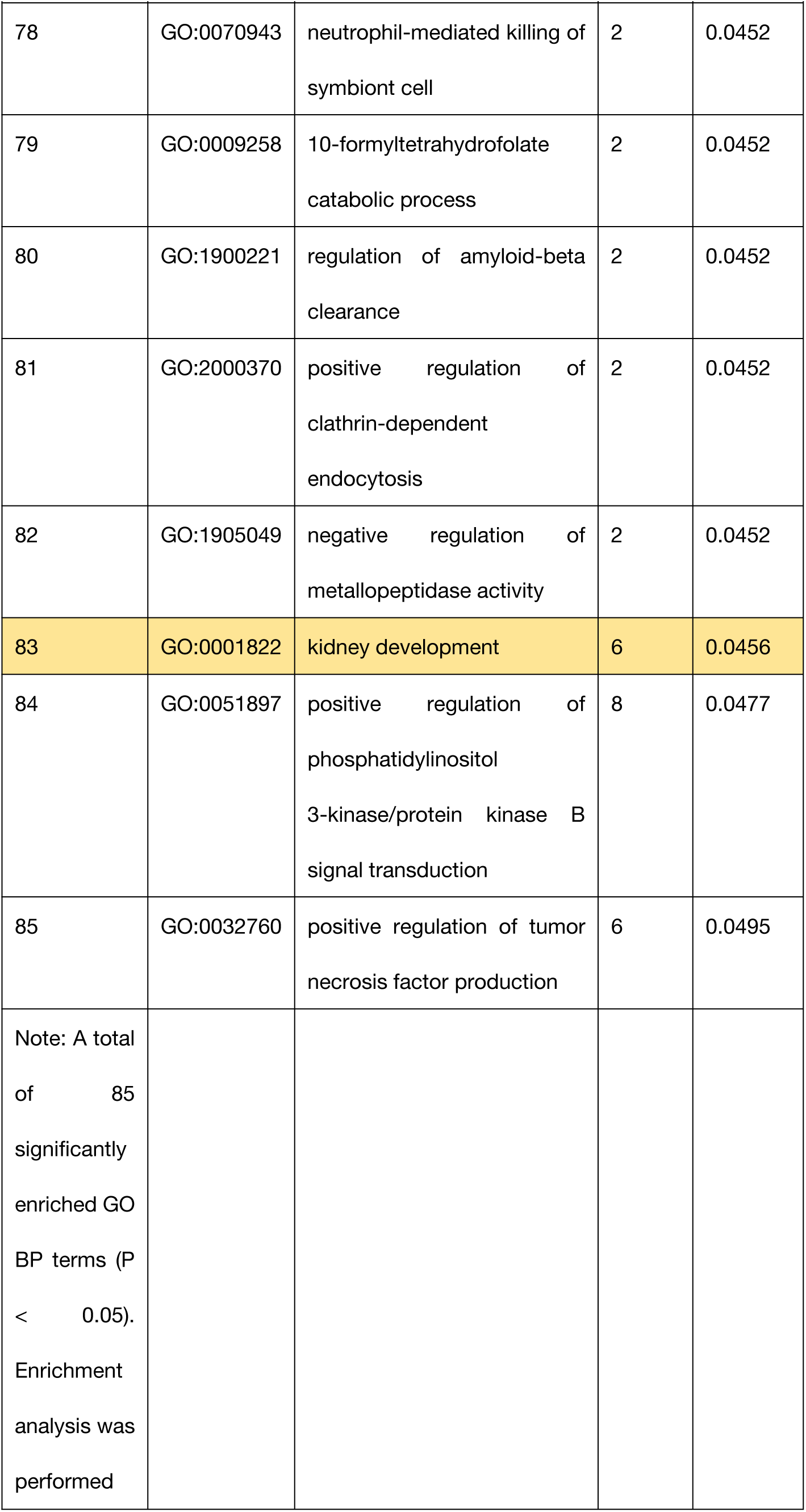
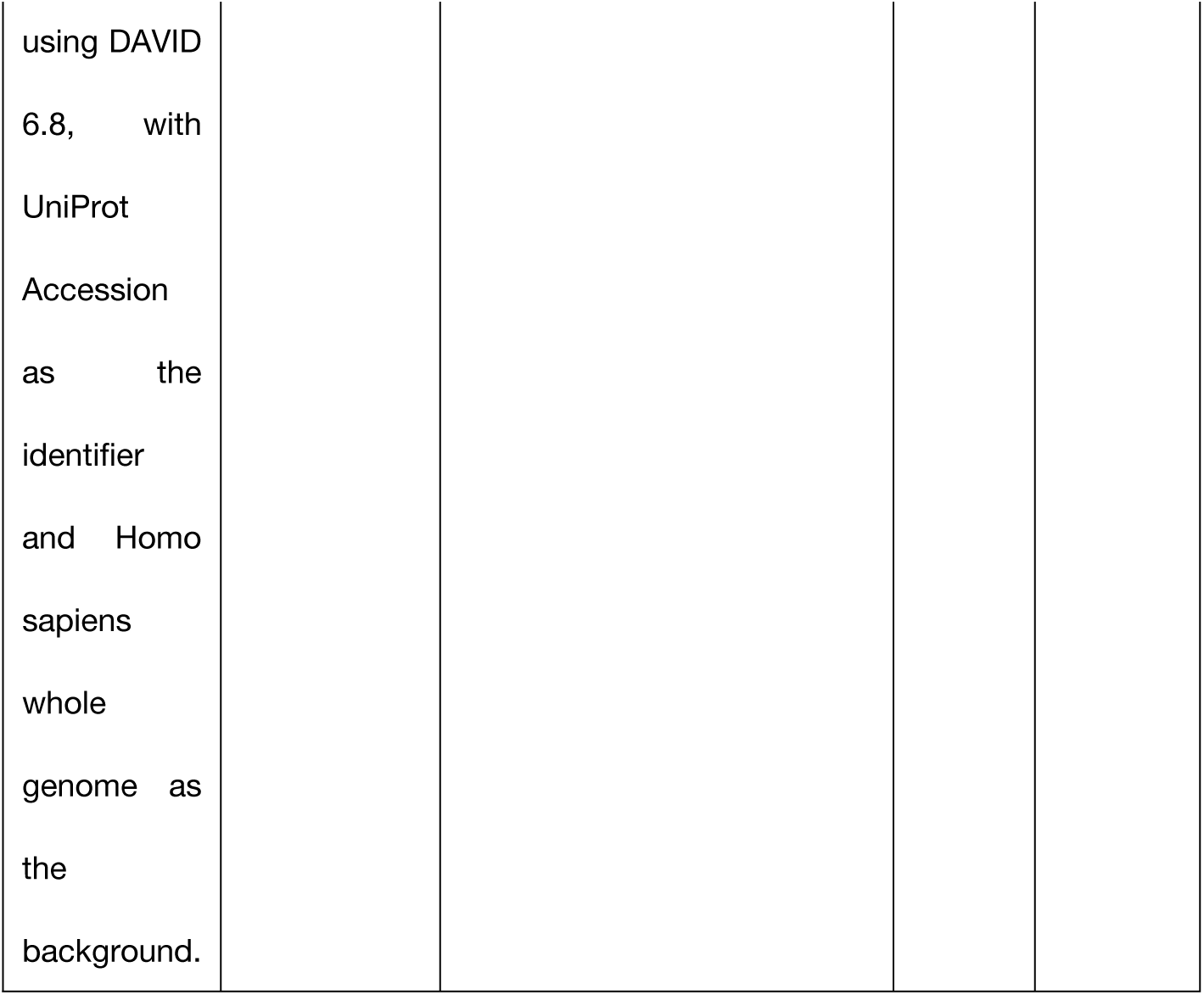
GO BP enrichment terms for R6 at 6–8 weeks of gestation (P < 0.05, 85 terms in total).

Subject R6 identified a total of 85 significantly enriched GO BP terms at 6–8 weeks of gestation (Table 2). Among them, 10 terms were related to organ/tissue development, including: spinal cord development (P = 1.16×10⁻²), angiogenesis (P = 2.77×10⁻²), loop of Henle development (P = 3.03×10⁻²), synapse assembly (P = 2.33×10⁻²), neuron projection development (P = 2.56×10⁻²), central nervous system development (P = 2.71×10⁻²), inner ear auditory receptor cell differentiation (P = 1.88×10⁻²), peripheral nervous system myelination (P = 4.04×10⁻²), and kidney development (P = 4.56×10⁻²).

Subject R15 identified a total of 51 significantly enriched GO BP terms at 6–8 weeks of gestation (Table 3). The organ/tissue development-related term was ossification (P = 3.20×10⁻²).

**Table 3.** GO BP enrichment terms for R15 at 6–8 weeks of gestation (P < 0.05, 51 terms in total).

| No. | GO ID | Biological Process<br>(English) | Count | P-value |
| --- | --- | --- | --- | --- |
| 1 | GO:0051764 | actin crosslink formation | 4 | 1.88e-04 |
| 2 | GO:0042542 | response to hydrogen peroxide | 5 | 2.21e-04 |
| 3 | GO:0051017 | actin filament bundle assembly | 5 | 2.21e-04 |
| 4 | GO:0032956 | regulation of actin cytoskeleton organization | 6 | 0.0014 |
| 5 | GO:0072659 | protein localization to plasma membrane | 7 | 0.0015 |
| 6 | GO:0016477 | cell migration | 9 | 0.0015 |
| 7 | GO:0007266 | Rho protein signal transduction | 5 | 0.0018 |
| 8 | GO:0006979 | response to oxidative stress | 7 | 0.0024 |
| 9 | GO:0045109 | intermediate filament organization | 5 | 0.0024 |
| 10 | GO:0031638 | zymogen activation | 4 | 0.0026 |
| 11 | GO:0046034 | ATP metabolic process | 4 | 0.0031 |
| 12 | GO:0006508 | proteolysis | 13 | 0.0041 |
| 13 | GO:0016236 | macroautophagy | 5 | 0.0043 |
| 14 | GO:0006958 | complement activation, classical pathway | 4 | 0.0043 |
| 15 | GO:0160257 | complement activation, GZMK pathway | 3 | 0.0050 |
| 16 | GO:0050778 | positive regulation of immune response | 4 | 0.0056 |
| 17 | GO:0070527 | platelet aggregation | 4 | 0.0059 |
| 18 | GO:0051897 | positive regulation of phosphatidylinositol<br>3-kinase/protein kinase B<br>signal transduction | 7 | 0.0063 |
| 19 | GO:1903543 | positive regulation of<br>exosomal secretion | 3 | 0.0065 |
| 20 | GO:0006957 | complement activation,<br>alternative pathway | 3 | 0.0092 |
| 21 | GO:0051016 | barbed-end actin filament<br>capping | 3 | 0.0101 |
| 22 | GO:0007009 | plasma membrane<br>organization | 3 | 0.0101 |
| 23 | GO:0046686 | response to cadmium ion | 3 | 0.0122 |
| 24 | GO:0008104 | intracellular protein<br>localization | 6 | 0.0126 |
| 25 | GO:0031424 | keratinization | 4 | 0.0179 |
| 26 | GO:0031100 | animal organ regeneration | 3 | 0.0207 |
| 27 | GO:0061621 | canonical glycolysis | 3 | 0.0207 |
| 28 | GO:0008360 | regulation of cell shape | 5 | 0.0208 |
| 29 | GO:0006858 | extracellular transport | 2 | 0.0228 |
| 30 | GO:2000296 | negative regulation of<br>hydrogen peroxide<br>catabolic process | 2 | 0.0228 |
| 31 | GO:1902903 | regulation of<br>supramolecular fiber<br>organization | 2 | 0.0228 |
| 32 | GO:1903551 | regulation of extracellular<br>exosome assembly | 2 | 0.0228 |
| 33 | GO:0070374 | positive regulation of ERK1<br>and ERK2 cascade | 6 | 0.0230 |
| 34 | GO:0006956 | complement activation | 3 | 0.0264 |
| 35 | GO:0036258 | multivesicular body<br>assembly | 3 | 0.0264 |
| 36 | GO:0006954 | inflammatory response | 9 | 0.0274 |
| 37 | GO:0032355 | response to estradiol | 4 | 0.0276 |
| 38 | GO:0000398 | mRNA splicing, via<br>spliceosome | 6 | 0.0293 |
| 39 | GO:0045732 | positive regulation of<br>protein catabolic process | 4 | 0.0302 |
| 40 | GO:1903553 | positive regulation of<br>extracellular exosome<br>assembly | 2 | 0.0303 |
| 41 | GO:0033591 | response to L-ascorbic<br>acid | 2 | 0.0303 |
| 42 | GO:0098609 | cell-cell adhesion | 6 | 0.0308 |
| 43 | GO:0001503 | ossification | 4 | 0.0320 |
| 44 | GO:2001237 | negative regulation of<br>extrinsic apoptotic<br>signaling pathway | 3 | 0.0360 |
| 45 | GO:1902115 | regulation of organelle<br>assembly | 2 | 0.0378 |
| 46 | GO:0010628 | positive regulation of gene<br>expression | 9 | 0.0387 |
| 47 | GO:0006953 | acute-phase response | 3 | 0.0394 |
| 48 | GO:0051546 | keratinocyte migration | 2 | 0.0451 |
| 49 | GO:0072318 | clathrin coat disassembly | 2 | 0.0451 |
| 50 | GO:0022617 | extracellular matrix<br>disassembly | 3 | 0.0466 |
| 51 | GO:0034976 | response to endoplasmic<br>reticulum stress | 4 | 0.0470 |
| Note: A total of 51 significantly enriched GO BP terms ($P < 0.05$). Enrichment analysis was performed using DAVID 6.8, with UniProt Accession as the identifier and Homo sapiens whole genome as the background. |  |  |  |  |

Subject R16 identified a total of 40 significantly enriched GO BP terms at 6–8 weeks of gestation (Table 4). The organ/tissue development-related term was keratinocyte differentiation (P = 1.15×10⁻²).

**Table 4.** GO BP enrichment terms for R16 at 6–8 weeks of gestation (P < 0.05, 40 terms in total).

| No. | GO ID | Biological Process (English) | Protein Count | P-value |
| --- | --- | --- | --- | --- |
| 1 | GO:0002227 | innate immune response in mucosa | 15 | 5.39e-22 |
| 2 | GO:0019731 | antibacterial humoral response | 15 | 2.65e-16 |
| 3 | GO:0061844 | antimicrobial humoral immune response mediated by antimicrobial peptide | 17 | 1.52e-15 |
| 4 | GO:0050830 | defense response to Gram-positive bacterium | 14 | 2.10e-11 |
| 5 | GO:0006325 | chromatin organization | 16 | 1.59e-08 |
| 6 | GO:0006334 | nucleosome assembly | 11 | 1.81e-07 |
| 7 | GO:0070527 | platelet aggregation | 6 | 1.94e-05 |
| 8 | GO:0031640 | killing of cells of another organism | 6 | 5.31e-04 |
| 9 | GO:0098869 | cellular oxidant detoxification | 5 | 9.34e-04 |
| 10 | GO:0061621 | canonical glycolysis | 4 | 0.0010 |
| 11 | GO:0034599 | cellular response to oxidative stress | 6 | 0.0012 |
| 12 | GO:1904874 | positive regulation of telomerase RNA localization to Cajal body | 3 | 0.0016 |
| 13 | GO:1904871 | positive regulation of protein localization to Cajal body | 3 | 0.0020 |
| 14 | GO:0050821 | protein stabilization | 8 | 0.0022 |
| 15 | GO:0006457 | protein folding | 7 | 0.0031 |
| 16 | GO:0098761 | cellular response to interleukin-7 | 3 | 0.0041 |
| 17 | GO:0031639 | plasminogen activation | 3 | 0.0041 |
| 18 | GO:1900026 | positive regulation of substrate adhesion-dependent cell spreading | 4 | 0.0042 |
| 19 | GO:0006096 | glycolytic process | 4 | 0.0052 |
| 20 | GO:0006094 | gluconeogenesis | 4 | 0.0065 |
| 21 | GO:0018149 | peptide cross-linking | 3 | 0.0067 |
| 22 | GO:0045104 | intermediate filament cytoskeleton organization | 3 | 0.0075 |
| 23 | GO:0008285 | negative regulation of cell population proliferation | 9 | 0.0102 |
| 24 | GO:0032212 | positive regulation of telomere maintenance via telomerase | 3 | 0.0108 |
| 25 | GO:0030216 | keratinocyte differentiation | 4 | 0.0115 |
| 26 | GO:0045109 | intermediate filament organization | 4 | 0.0140 |
| 27 | GO:0030451 | regulation of complement activation, alternative pathway | 2 | 0.0205 |
| 28 | GO:0001970 | positive regulation of activation of membrane attack complex | 2 | 0.0205 |
| 29 | GO:0006956 | complement activation | 3 | 0.0216 |
| 30 | GO:0050832 | defense response to fungus | 3 | 0.0241 |
| 31 | GO:0007165 | signal transduction | 17 | 0.0294 |
| 32 | GO:2000352 | negative regulation of<br>endothelial cell apoptotic<br>process | 3 | 0.0295 |
| 33 | GO:0007339 | binding of sperm to zona<br>pellucida | 3 | 0.0339 |
| 34 | GO:0090666 | scaRNA localization to Cajal<br>body | 2 | 0.0340 |
| 35 | GO:0045454 | cell redox homeostasis | 3 | 0.0416 |
| 36 | GO:0050714 | positive regulation of protein<br>secretion | 3 | 0.0448 |
| 37 | GO:0071364 | cellular response to<br>epidermal growth factor<br>stimulus | 3 | 0.0448 |
| 38 | GO:0099638 | endosome to plasma<br>membrane protein transport | 2 | 0.0472 |
| 39 | GO:0051673 | disruption of plasma<br>membrane integrity in<br>another organism | 2 | 0.0472 |
| 40 | GO:0006166 | purine ribonucleoside<br>salvage | 2 | 0.0472 |
| Note: A total<br>of 40<br>significantly<br>enriched GO<br>BP terms (P<br>< 0.05). |  |  |  |  |
| Enrichment analysis was performed using DAVID 6.8, with UniProt Accession as the identifier and Homo sapiens whole genome as the background. |  |  |  |  |

Subject R6 identified a total of 269 significantly enriched GO BP terms at 22–24 weeks of gestation (Table 5). R6-T2 was the sample with the largest number and richest signals among the nine samples. Among them, 51 terms were related to organ/tissue development and morphogenesis, covering multiple organ systems, including axon guidance (P = 3.36×10⁻⁹), angiogenesis (P = 4.71×10⁻⁷), nervous system development (P = 1.89×10⁻⁵), placenta blood vessel development (P = 2.45×10⁻⁴), intrahepatic bile duct development (P = 6.48×10⁻⁴), podocyte development (P = 1.07×10⁻³), pulmonary valve morphogenesis (P = 1.35×10⁻³), pericardium development (P = 3.30×10⁻³), embryonic limb morphogenesis (P = 2.22×10⁻³), animal organ morphogenesis (P = 2.72×10⁻³), animal organ development (P = 2.90×10⁻³), atrioventricular node development (P = 4.81×10⁻³), kidney development (P = 6.06×10⁻³), in utero embryonic development (P = 1.04×10⁻²), lung development (P = 2.33×10⁻²), heart development (P = 2.36×10⁻²), ossification (P = 3.15×10⁻²), and inner ear development (P = 4.97×10⁻²). Multiple organ-development-related terms had P-values below 10⁻³ and covered multiple organ systems including nervous, vascular, cardiac, renal, pulmonary, and auditory, suggesting that the urinary proteome captured abundant fetal organ development signals at this time point.

**Table 5.** GO BP enrichment terms for R6 at 22–24 weeks of gestation (P < 0.05, 269 terms in total).

| No. | GO ID | Biological Process<br>(English) | Count | P-value |
| --- | --- | --- | --- | --- |
| 1 | GO:0007155 | cell adhesion | 94 | 1.29e-28 |
| 2 | GO:0007165 | signal transduction | 108 | 1.51e-10 |
| 3 | GO:0007411 | axon guidance | 30 | 3.36e-09 |
| 4 | GO:0098609 | cell-cell adhesion | 32 | 4.27e-09 |
| 5 | GO:0007157 | heterophilic cell-cell adhesion | 16 | 4.68e-09 |
| 6 | GO:0007156 | homophilic cell-cell<br>adhesion | 28 | 5.54e-09 |
| 7 | GO:0016477 | cell migration | 36 | 6.15e-09 |
| 8 | GO:0034315 | regulation of Arp2/3<br>complex-mediated actin<br>nucleation | 11 | 9.03e-09 |
| 9 | GO:0030335 | positive regulation of cell<br>migration | 36 | 1.81e-08 |
| 10 | GO:0006897 | endocytosis | 29 | 3.64e-08 |
| 11 | GO:0007596 | blood coagulation | 20 | 4.07e-08 |
| 12 | GO:0006955 | immune response | 53 | 1.01e-07 |
| 13 | GO:0006898 | receptor-mediated<br>endocytosis | 17 | 1.85e-07 |
| 14 | GO:0007565 | female pregnancy | 17 | 1.85e-07 |
| 15 | GO:0030334 | regulation of cell migration | 20 | 3.26e-07 |
| 16 | GO:0001525 | angiogenesis | 32 | 4.71e-07 |
| 17 | GO:0036258 | multivesicular body<br>assembly | 11 | 5.03e-07 |
| 18 | GO:0007166 | cell surface receptor<br>signaling pathway | 39 | 8.41e-07 |
| 19 | GO:0007160 | cell-matrix adhesion | 19 | 8.91e-07 |
| 20 | GO:0031623 | receptor internalization | 13 | 9.57e-07 |
| 21 | GO:0015031 | protein transport | 49 | 9.94e-07 |
| 22 | GO:0009313 | oligosaccharide catabolic<br>process | 7 | 1.82e-06 |
| 23 | GO:0043410 | positive regulation of MAPK<br>cascade | 27 | 2.38e-06 |
| 24 | GO:0008284 | positive regulation of cell<br>population proliferation | 46 | 3.79e-06 |
| 25 | GO:0090148 | membrane fission | 11 | 4.52e-06 |
| 26 | GO:0030168 | platelet activation | 13 | 1.35e-05 |
| 27 | GO:0070374 | positive regulation of ERK1<br>and ERK2 cascade | 24 | 1.87e-05 |
| 28 | GO:0007399 | nervous system<br>development | 44 | 1.89e-05 |
| 29 | GO:0009653 | anatomical structure<br>morphogenesis | 20 | 2.93e-05 |
| 30 | GO:0007162 | negative regulation of cell<br>adhesion | 11 | 2.97e-05 |
| 31 | GO:0039702 | viral budding via host<br>ESCRT complex | 8 | 4.81e-05 |
| 32 | GO:0006954 | inflammatory response | 39 | 4.84e-05 |
| 33 | GO:0002250 | adaptive immune response | 41 | 5.38e-05 |
| 34 | GO:0022617 | extracellular matrix<br>disassembly | 10 | 7.41e-05 |
| 35 | GO:0051897 | positive regulation of<br>phosphatidylinositol<br>3-kinase/protein kinase B<br>signal transduction | 23 | 7.91e-05 |
| 36 | GO:0018108 | peptidyl-tyrosine<br>phosphorylation | 9 | 8.47e-05 |
| 37 | GO:0046579 | positive regulation of Ras<br>protein signal transduction | 8 | 1.06e-04 |
| 38 | GO:0030195 | negative regulation of blood coagulation | 6 | 1.69e-04 |
| 39 | GO:0007169 | cell surface receptor<br>protein tyrosine kinase<br>signaling pathway | 16 | 1.72e-04 |
| 40 | GO:0019221 | cytokine-mediated<br>signaling pathway | 18 | 1.87e-04 |
| 41 | GO:0016197 | endosomal transport | 13 | 2.21e-04 |
| 42 | GO:0090263 | positive regulation of<br>canonical Wnt signaling<br>pathway | 16 | 2.25e-04 |
| 43 | GO:0060674 | placenta blood vessel<br>development | 6 | 2.45e-04 |
| 44 | GO:0030155 | regulation of cell adhesion | 11 | 2.61e-04 |
| 45 | GO:0045672 | positive regulation of<br>osteoclast differentiation | 8 | 3.21e-04 |
| 46 | GO:0035633 | maintenance of blood-brain<br>barrier | 8 | 3.21e-04 |
| 47 | GO:1903543 | positive regulation of<br>exosomal secretion | 6 | 3.44e-04 |
| 48 | GO:0034446 | substrate<br>adhesion-dependent cell<br>spreading | 10 | 3.69e-04 |
| 49 | GO:0046718 | symbiont entry into host<br>cell | 11 | 3.82e-04 |
| 50 | GO:0006508 | proteolysis | 46 | 3.84e-04 |
| 51 | GO:0043162 | ubiquitin-dependent protein<br>catabolic process via the multivesicular body sorting<br>pathway | 8 | 3.90e-04 |
| 52 | GO:0099560 | synaptic membrane<br>adhesion | 8 | 3.90e-04 |
| 53 | GO:0051384 | response to glucocorticoid | 10 | 4.24e-04 |
| 54 | GO:0007042 | lysosomal lumen<br>acidification | 7 | 4.25e-04 |
| 55 | GO:0032956 | regulation of actin<br>cytoskeleton organization | 14 | 4.39e-04 |
| 56 | GO:0072574 | hepatocyte proliferation | 5 | 4.85e-04 |
| 57 | GO:0007267 | cell-cell signaling | 22 | 5.42e-04 |
| 58 | GO:0016339 | calcium-dependent cell-cell<br>adhesion | 9 | 5.91e-04 |
| 59 | GO:2001204 | regulation of osteoclast<br>development | 4 | 6.48e-04 |
| 60 | GO:0035622 | intrahepatic bile duct<br>development | 4 | 6.48e-04 |
| 61 | GO:0008038 | neuron recognition | 4 | 6.48e-04 |
| 62 | GO:1990705 | cholangiocyte proliferation | 4 | 6.48e-04 |
| 63 | GO:0070527 | platelet aggregation | 9 | 6.84e-04 |
| 64 | GO:0001755 | neural crest cell migration | 9 | 6.84e-04 |
| 65 | GO:0071526 | semaphorin-plexin<br>signaling pathway | 9 | 6.84e-04 |
| 66 | GO:0051894 | positive regulation of focal<br>adhesion assembly | 7 | 8.15e-04 |
| 67 | GO:0045216 | cell-cell junction<br>organization | 7 | 9.92e-04 |
| 68 | GO:0030163 | protein catabolic process | 15 | 0.0011 |
| 69 | GO:0046849 | bone remodeling | 6 | 0.0011 |
| 70 | GO:0072015 | podocyte development | 5 | 0.0011 |
| 71 | GO:0007229 | integrin-mediated signaling<br>pathway | 14 | 0.0011 |
| 72 | GO:0045954 | positive regulation of<br>natural killer cell mediated<br>cytotoxicity | 7 | 0.0012 |
| 73 | GO:0110011 | regulation of basement<br>membrane organization | 4 | 0.0013 |
| 74 | GO:0072104 | glomerular capillary<br>formation | 4 | 0.0013 |
| 75 | GO:0031507 | heterochromatin formation | 15 | 0.0013 |
| 76 | GO:0003184 | pulmonary valve<br>morphogenesis | 6 | 0.0014 |
| 77 | GO:0008360 | regulation of cell shape | 15 | 0.0014 |
| 78 | GO:0005975 | carbohydrate metabolic<br>process | 18 | 0.0015 |
| 79 | GO:0002011 | morphogenesis of an<br>epithelial sheet | 5 | 0.0015 |
| 80 | GO:2001046 | positive regulation of<br>integrin-mediated signaling<br>pathway | 5 | 0.0015 |
| 81 | GO:0050679 | positive regulation of<br>epithelial cell proliferation | 10 | 0.0016 |
| 82 | GO:0030501 | positive regulation of bone<br>mineralization | 8 | 0.0017 |
| 83 | GO:0019886 | antigen processing and presentation of exogenous peptide antigen via MHC class II | 7 | 0.0017 |
| 84 | GO:0048146 | positive regulation of fibroblast proliferation | 9 | 0.0017 |
| 85 | GO:0016192 | vesicle-mediated transport | 26 | 0.0018 |
| 86 | GO:0071228 | cellular response to tumor cell | 4 | 0.0021 |
| 87 | GO:0061073 | ciliary body morphogenesis | 4 | 0.0021 |
| 88 | GO:0030326 | embryonic limb morphogenesis | 8 | 0.0022 |
| 89 | GO:0030198 | extracellular matrix organization | 16 | 0.0023 |
| 90 | GO:0010811 | positive regulation of cell-substrate adhesion | 7 | 0.0024 |
| 91 | GO:0001934 | positive regulation of protein phosphorylation | 9 | 0.0024 |
| 92 | GO:0006886 | intracellular protein transport | 24 | 0.0024 |
| 93 | GO:0019882 | antigen processing and presentation | 8 | 0.0025 |
| 94 | GO:0016236 | macroautophagy | 11 | 0.0027 |
| 95 | GO:0007566 | embryo implantation | 8 | 0.0029 |
| 96 | GO:0048513 | animal organ development | 13 | 0.0029 |
| 97 | GO:0045785 | positive regulation of cell adhesion | 10 | 0.0029 |
| 98 | GO:0071985 | multivesicular body sorting pathway | 6 | 0.0031 |
| 99 | GO:0031638 | zymogen activation | 7 | 0.0032 |
| 100 | GO:0006689 | ganglioside catabolic process | 4 | 0.0033 |
| 101 | GO:0060039 | pericardium development | 4 | 0.0033 |
| 102 | GO:0072014 | proximal tubule development | 4 | 0.0033 |
| 103 | GO:0043328 | protein transport to vacuole involved in ubiquitin-dependent protein catabolic process via the multivesicular body sorting pathway | 5 | 0.0034 |
| 104 | GO:0051149 | positive regulation of muscle cell differentiation | 5 | 0.0034 |
| 105 | GO:0010824 | regulation of centrosome duplication | 6 | 0.0037 |
| 106 | GO:0042147 | retrograde transport, endosome to Golgi | 11 | 0.0041 |
| 107 | GO:0006044 | N-acetylglucosamine metabolic process | 5 | 0.0044 |
| 108 | GO:0070986 | left/right axis specification | 5 | 0.0044 |
| 109 | GO:0061952 | midbody abscission | 6 | 0.0044 |
| 110 | GO:0042060 | wound healing | 10 | 0.0046 |
| 111 | GO:0001649 | osteoblast differentiation | 14 | 0.0048 |
| 112 | GO:0010544 | negative regulation of platelet activation | 4 | 0.0048 |
| 113 | GO:0003162 | atrioventricular node development | 4 | 0.0048 |
| 114 | GO:1900426 | positive regulation of defense response to bacterium | 4 | 0.0048 |
| 115 | GO:0031667 | response to nutrient levels | 10 | 0.0050 |
| 116 | GO:0002376 | immune system process | 14 | 0.0051 |
| 117 | GO:0016064 | immunoglobulin mediated immune response | 12 | 0.0052 |
| 118 | GO:0071346 | cellular response to type II interferon | 10 | 0.0054 |
| 119 | GO:0044331 | cell-cell adhesion mediated by cadherin | 7 | 0.0055 |
| 120 | GO:0007259 | cell surface receptor signaling pathway via JAK-STAT | 10 | 0.0059 |
| 121 | GO:0001822 | kidney development | 13 | 0.0061 |
| 122 | GO:0050793 | regulation of developmental process | 7 | 0.0062 |
| 123 | GO:0002682 | regulation of immune system process | 8 | 0.0063 |
| 124 | GO:0007416 | synapse assembly | 10 | 0.0064 |
| 125 | GO:0050830 | defense response to Gram-positive bacterium | 14 | 0.0064 |
| 126 | GO:1904903 | ESCRT III complex disassembly | 4 | 0.0067 |
| 127 | GO:1901203 | positive regulation of extracellular matrix assembly | 4 | 0.0067 |
| 128 | GO:0030574 | collagen catabolic process | 7 | 0.0070 |
| 129 | GO:0016601 | Rac protein signal transduction | 6 | 0.0070 |
| 130 | GO:0051592 | response to calcium ion | 8 | 0.0077 |
| 131 | GO:0051496 | positive regulation of stress fiber assembly | 8 | 0.0077 |
| 132 | GO:0030513 | positive regulation of BMP signaling pathway | 7 | 0.0078 |
| 133 | GO:0048010 | vascular endothelial growth factor receptor signaling pathway | 6 | 0.0080 |
| 134 | GO:0007413 | axonal fasciculation | 5 | 0.0081 |
| 135 | GO:0050808 | synapse organization | 9 | 0.0084 |
| 136 | GO:0043524 | negative regulation of neuron apoptotic process | 15 | 0.0086 |
| 137 | GO:0032509 | endosome transport via multivesicular body sorting pathway | 4 | 0.0089 |
| 138 | GO:0045428 | regulation of nitric oxide biosynthetic process | 4 | 0.0089 |
| 139 | GO:0042117 | monocyte activation | 4 | 0.0089 |
| 140 | GO:0003140 | determination of left/right asymmetry in lateral mesoderm | 4 | 0.0089 |
| 141 | GO:0030097 | hemopoiesis | 9 | 0.0092 |
| 142 | GO:0019082 | viral protein processing | 6 | 0.0092 |
| 143 | GO:0045995 | regulation of embryonic development | 8 | 0.0093 |
| 144 | GO:1905521 | regulation of macrophage migration | 3 | 0.0096 |
| 145 | GO:0030209 | dermatan sulfate proteoglycan catabolic process | 3 | 0.0096 |
| 146 | GO:1903553 | positive regulation of extracellular exosome assembly | 3 | 0.0096 |
| 147 | GO:0099537 | trans-synaptic signaling | 3 | 0.0096 |
| 148 | GO:0060666 | dichotomous subdivision of terminal units involved in salivary gland branching | 3 | 0.0096 |
| 149 | GO:1990182 | exosomal secretion | 3 | 0.0096 |
| 150 | GO:0043011 | myeloid dendritic cell differentiation | 5 | 0.0097 |
| 151 | GO:0031468 | nuclear membrane reassembly | 5 | 0.0097 |
| 152 | GO:0002437 | inflammatory response to antigenic stimulus | 5 | 0.0097 |
| 153 | GO:0050776 | regulation of immune response | 7 | 0.0098 |
| 154 | GO:0001701 | in utero embryonic development | 19 | 0.0104 |
| 155 | GO:0070372 | regulation of ERK1 and ERK2 cascade | 6 | 0.0105 |
| 156 | GO:0034332 | adherens junction organization | 6 | 0.0105 |
| 157 | GO:0042177 | negative regulation of protein catabolic process | 7 | 0.0108 |
| 158 | GO:0045087 | innate immune response | 36 | 0.0114 |
| 159 | GO:0045807 | positive regulation of endocytosis | 5 | 0.0114 |
| 160 | GO:0010744 | positive regulation of macrophage derived foam cell differentiation | 5 | 0.0114 |
| 161 | GO:0051918 | negative regulation of fibrinolysis | 4 | 0.0115 |
| 162 | GO:0002315 | marginal zone B cell differentiation | 4 | 0.0115 |
| 163 | GO:0048251 | elastic fiber assembly | 4 | 0.0115 |
| 164 | GO:1900026 | positive regulation of substrate adhesion-dependent cell spreading | 7 | 0.0120 |
| 165 | GO:1902533 | positive regulation of intracellular signal transduction | 7 | 0.0120 |
| 166 | GO:1902895 | positive regulation of miRNA transcription | 8 | 0.0133 |
| 167 | GO:0008045 | motor neuron axon guidance | 5 | 0.0134 |
| 168 | GO:0007034 | vacuolar transport | 5 | 0.0134 |
| 169 | GO:0051591 | response to cAMP | 6 | 0.0135 |
| 170 | GO:0032729 | positive regulation of type II<br>interferon production | 10 | 0.0141 |
| 171 | GO:0010572 | positive regulation of<br>platelet activation | 4 | 0.0145 |
| 172 | GO:0032722 | positive regulation of<br>chemokine production | 7 | 0.0146 |
| 173 | GO:0072659 | protein localization to<br>plasma membrane | 14 | 0.0148 |
| 174 | GO:0006935 | chemotaxis | 13 | 0.0149 |
| 175 | GO:0034197 | triglyceride transport | 3 | 0.0155 |
| 176 | GO:0071726 | cellular response to diacyl<br>bacterial lipopeptide | 3 | 0.0155 |
| 177 | GO:0017015 | regulation of transforming<br>growth factor beta receptor<br>signaling pathway | 5 | 0.0156 |
| 178 | GO:0031333 | negative regulation of<br>protein-containing complex<br>assembly | 6 | 0.0169 |
| 179 | GO:0007159 | leukocyte cell-cell adhesion | 6 | 0.0169 |
| 180 | GO:0051152 | positive regulation of<br>smooth muscle cell<br>differentiation | 4 | 0.0179 |
| 181 | GO:2001028 | positive regulation of<br>endothelial cell chemotaxis | 4 | 0.0179 |
| 182 | GO:0034314 | Arp2/3 complex-mediated<br>actin nucleation | 5 | 0.0179 |
| 183 | GO:0030301 | cholesterol transport | 5 | 0.0179 |
| 184 | GO:0031175 | neuron projection<br>development | 13 | 0.0181 |
| 185 | GO:0032760 | positive regulation of tumor<br>necrosis factor production | 12 | 0.0182 |
| 186 | GO:0051963 | regulation of synapse<br>assembly | 6 | 0.0188 |
| 187 | GO:0042127 | regulation of cell population<br>proliferation | 15 | 0.0197 |
| 188 | GO:0016525 | negative regulation of<br>angiogenesis | 12 | 0.0211 |
| 189 | GO:0001947 | heart looping | 8 | 0.0212 |
| 190 | GO:0010629 | negative regulation of gene<br>expression | 23 | 0.0215 |
| 191 | GO:0060396 | growth hormone receptor<br>signaling pathway | 4 | 0.0217 |
| 192 | GO:0030194 | positive regulation of blood<br>coagulation | 4 | 0.0217 |
| 193 | GO:0045651 | positive regulation of<br>macrophage differentiation | 4 | 0.0217 |
| 194 | GO:0006750 | glutathione biosynthetic<br>process | 4 | 0.0217 |
| 195 | GO:0097061 | dendritic spine organization | 3 | 0.0226 |
| 196 | GO:0002860 | positive regulation of<br>natural killer cell mediated cytotoxicity directed<br>against tumor cell target | 3 | 0.0226 |
| 197 | GO:0006572 | L-tyrosine catabolic<br>process | 3 | 0.0226 |
| 198 | GO:1900108 | negative regulation of nodal<br>signaling pathway | 3 | 0.0226 |
| 199 | GO:0090611 | ubiquitin-independent<br>protein catabolic process<br>via the multivesicular body<br>sorting pathway | 3 | 0.0226 |
| 200 | GO:1902396 | protein localization to<br>bicellular tight junction | 3 | 0.0226 |
| 201 | GO:0050919 | negative chemotaxis | 6 | 0.0231 |
| 202 | GO:0030324 | lung development | 9 | 0.0233 |
| 203 | GO:0006911 | phagocytosis, engulfment | 5 | 0.0233 |
| 204 | GO:0002548 | monocyte chemotaxis | 5 | 0.0233 |
| 205 | GO:0007507 | heart development | 17 | 0.0236 |
| 206 | GO:0060413 | atrial septum<br>morphogenesis | 4 | 0.0259 |
| 207 | GO:0099558 | maintenance of synapse<br>structure | 4 | 0.0259 |
| 208 | GO:0006796 | phosphate-containing<br>compound metabolic<br>process | 5 | 0.0264 |
| 209 | GO:1901673 | regulation of mitotic spindle<br>assembly | 5 | 0.0264 |
| 210 | GO:0001954 | positive regulation of cell-matrix adhesion | 5 | 0.0264 |
| 211 | GO:0048545 | response to steroid hormone | 5 | 0.0264 |
| 212 | GO:0009887 | animal organ morphogenesis | 10 | 0.0272 |
| 213 | GO:0010718 | positive regulation of epithelial to mesenchymal transition | 7 | 0.0289 |
| 214 | GO:0017157 | regulation of exocytosis | 5 | 0.0296 |
| 215 | GO:0032456 | endocytic recycling | 9 | 0.0297 |
| 216 | GO:0030214 | hyaluronan catabolic process | 4 | 0.0305 |
| 217 | GO:0099151 | regulation of postsynaptic density assembly | 4 | 0.0305 |
| 218 | GO:0030031 | cell projection assembly | 4 | 0.0305 |
| 219 | GO:0042756 | drinking behavior | 4 | 0.0305 |
| 220 | GO:0046761 | viral budding from plasma membrane | 4 | 0.0305 |
| 221 | GO:0032482 | Rab protein signal transduction | 4 | 0.0305 |
| 222 | GO:0060391 | positive regulation of SMAD protein signal transduction | 6 | 0.0306 |
| 223 | GO:0009967 | positive regulation of signal transduction | 6 | 0.0306 |
| 224 | GO:0061450 | trophoblast cell migration | 3 | 0.0308 |
| 225 | GO:0023056 | positive regulation of signaling | 3 | 0.0308 |
| 226 | GO:0071635 | negative regulation of transforming growth factor beta production | 3 | 0.0308 |
| 227 | GO:0001503 | ossification | 9 | 0.0315 |
| 228 | GO:0043065 | positive regulation of apoptotic process | 22 | 0.0316 |
| 229 | GO:0010759 | positive regulation of macrophage chemotaxis | 5 | 0.0331 |
| 230 | GO:0042832 | defense response to protozoan | 5 | 0.0331 |
| 231 | GO:0031295 | T cell costimulation | 6 | 0.0334 |
| 232 | GO:0007032 | endosome organization | 6 | 0.0334 |
| 233 | GO:1903078 | positive regulation of protein localization to plasma membrane | 7 | 0.0335 |
| 234 | GO:0006887 | exocytosis | 11 | 0.0347 |
| 235 | GO:0050852 | T cell receptor signaling pathway | 11 | 0.0364 |
| 236 | GO:0006997 | nucleus organization | 5 | 0.0368 |
| 237 | GO:0042531 | positive regulation of tyrosine phosphorylation of STAT protein | 5 | 0.0368 |
| 238 | GO:0006909 | phagocytosis | 8 | 0.0385 |
| 239 | GO:0010628 | positive regulation of gene expression | 30 | 0.0388 |
| 240 | GO:0032526 | response to retinoic acid | 6 | 0.0394 |
| 241 | GO:0006559 | L-phenylalanine catabolic process | 3 | 0.0400 |
| 242 | GO:0046755 | viral budding | 3 | 0.0400 |
| 243 | GO:0150094 | amyloid-beta clearance by<br>cellular catabolic process | 3 | 0.0400 |
| 244 | GO:0034372 | very-low-density lipoprotein<br>particle remodeling | 3 | 0.0400 |
| 245 | GO:0010647 | positive regulation of cell<br>communication | 3 | 0.0400 |
| 246 | GO:1902107 | positive regulation of<br>leukocyte differentiation | 3 | 0.0400 |
| 247 | GO:0035771 | interleukin-4-mediated<br>signaling pathway | 3 | 0.0400 |
| 248 | GO:1900738 | positive regulation of<br>phospholipase C-activating<br>G protein-coupled receptor<br>signaling pathway | 3 | 0.0400 |
| 249 | GO:0009896 | positive regulation of<br>catabolic process | 3 | 0.0400 |
| 250 | GO:0002090 | regulation of receptor<br>internalization | 3 | 0.0400 |
| 251 | GO:0001553 | luteinization | 3 | 0.0400 |
| 252 | GO:0050819 | negative regulation of<br>coagulation | 3 | 0.0400 |
| 253 | GO:0060346 | bone trabecula formation | 3 | 0.0400 |
| 254 | GO:0043066 | negative regulation of<br>apoptotic process | 32 | 0.0402 |
| 255 | GO:0035025 | positive regulation of Rho protein signal transduction | 5 | 0.0407 |
| 256 | GO:0010669 | epithelial structure maintenance | 4 | 0.0409 |
| 257 | GO:0097401 | synaptic vesicle lumen acidification | 4 | 0.0409 |
| 258 | GO:0006888 | endoplasmic reticulum to Golgi vesicle-mediated transport | 11 | 0.0416 |
| 259 | GO:0046427 | positive regulation of receptor signaling pathway via JAK-STAT | 6 | 0.0427 |
| 260 | GO:0030001 | metal ion transport | 5 | 0.0449 |
| 261 | GO:0002767 | immune response-inhibiting cell surface receptor signaling pathway | 5 | 0.0449 |
| 262 | GO:0006869 | lipid transport | 10 | 0.0450 |
| 263 | GO:0006091 | generation of precursor metabolites and energy | 6 | 0.0461 |
| 264 | GO:0050901 | leukocyte tethering or rolling | 4 | 0.0467 |
| 265 | GO:0022407 | regulation of cell-cell adhesion | 4 | 0.0467 |
| 266 | GO:0038084 | vascular endothelial growth factor signaling pathway | 4 | 0.0467 |
| 267 | GO:0002764 | immune response-regulating signaling pathway | 7 | 0.0470 |
| 268 | GO:0034599 | cellular response to oxidative stress | 10 | 0.0471 |
| 269 | GO:0048839 | inner ear development | 6 | 0.0497 |
| Note: A total of 269 significantly enriched GO BP terms ($P < 0.05$). Enrichment analysis was performed using DAVID 6.8, with UniProt Accession as the identifier and Homo sapiens whole genome as the background. |  |  |  |  |

Subject R15 identified a total of 95 significantly enriched GO BP terms at 22–24 weeks of gestation (Table 6). The organ/tissue development-related terms included: epidermis development (P = 1.21×10⁻⁴), angiogenesis (P = 1.48×10⁻²), cerebellar Purkinje cell layer development (P = 2.63×10⁻²), establishment of left/right asymmetry in lateral mesoderm (P = 2.63×10⁻²), and regulation of embryonic development (P = 1.15×10⁻²). In addition, chromatin remodeling terms were highly enriched, the most significant being telomere organization (P = 3.06×10⁻³⁴) and protein localization to CENP-A-containing chromatin (P = 4.22×10⁻¹⁸).

**Table 6.** GO BP enrichment terms for R15 at 22–24 weeks of gestation (P < 0.05, 95 terms in total).

| No. | GO ID | Biological Process<br>(English) | Count | P-value |
| --- | --- | --- | --- | --- |
| 1 | GO:0032200 | telomere organization | 24 | 3.06e-34 |
| 2 | GO:0061644 | protein localization to<br>CENP-A containing<br>chromatin | 14 | 4.22e-18 |
| 3 | GO:0045653 | negative regulation of<br>megakaryocyte<br>differentiation | 14 | 3.65e-17 |
| 4 | GO:0006334 | nucleosome assembly | 25 | 6.17e-13 |
| 5 | GO:0040029 | epigenetic regulation of gene expression | 15 | 4.89e-10 |
| 6 | GO:0006325 | chromatin organization | 26 | 6.97e-07 |
| 7 | GO:0008544 | epidermis development | 11 | 1.21e-04 |
| 8 | GO:0007155 | cell adhesion | 31 | 1.28e-04 |
| 9 | GO:0098609 | cell-cell adhesion | 16 | 3.26e-04 |
| 10 | GO:0001921 | positive regulation of receptor recycling | 5 | 3.33e-04 |
| 11 | GO:0006414 | translational elongation | 5 | 7.06e-04 |
| 12 | GO:0007015 | actin filament organization | 12 | 0.0014 |
| 13 | GO:0006886 | intracellular protein transport | 17 | 0.0017 |
| 14 | GO:0016192 | vesicle-mediated transport | 18 | 0.0017 |
| 15 | GO:0002768 | immune response-regulating cell surface receptor signaling pathway | 4 | 0.0024 |
| 16 | GO:0045063 | T-helper 1 cell differentiation | 5 | 0.0039 |
| 17 | GO:0017157 | regulation of exocytosis | 5 | 0.0045 |
| 18 | GO:0031507 | heterochromatin formation | 10 | 0.0046 |
| 19 | GO:0032434 | regulation of proteasomal ubiquitin-dependent protein catabolic process | 4 | 0.0047 |
| 20 | GO:0160165 | CD8-positive, alpha-beta T cell homeostasis | 4 | 0.0057 |
| 21 | GO:0043374 | CD8-positive, alpha-beta T cell differentiation | 4 | 0.0068 |
| 22 | GO:0045061 | thymic T cell selection | 4 | 0.0068 |
| 23 | GO:0031424 | keratinization | 7 | 0.0069 |
| 24 | GO:0045785 | positive regulation of cell<br>adhesion | 7 | 0.0073 |
| 25 | GO:0010498 | proteasomal protein<br>catabolic process | 7 | 0.0073 |
| 26 | GO:1900108 | negative regulation of nodal<br>signaling pathway | 3 | 0.0077 |
| 27 | GO:0070542 | response to fatty acid | 4 | 0.0081 |
| 28 | GO:0018149 | peptide cross-linking | 4 | 0.0081 |
| 29 | GO:0006979 | response to oxidative stress | 11 | 0.0088 |
| 30 | GO:0072539 | T-helper 17 cell<br>differentiation | 4 | 0.0094 |
| 31 | GO:0010499 | proteasomal<br>ubiquitin-independent<br>protein catabolic process | 4 | 0.0094 |
| 32 | GO:0048678 | response to axon injury | 5 | 0.0097 |
| 33 | GO:0060828 | regulation of canonical Wnt<br>signaling pathway | 5 | 0.0097 |
| 34 | GO:0006887 | exocytosis | 9 | 0.0098 |
| 35 | GO:0006955 | immune response | 24 | 0.0100 |
| 36 | GO:0007156 | homophilic cell-cell adhesion | 11 | 0.0102 |
| 37 | GO:0070358 | actin<br>polymerization-dependent<br>cell motility | 3 | 0.0107 |
| 38 | GO:0006405 | RNA export from nucleus | 4 | 0.0109 |
| 39 | GO:0045995 | regulation of embryonic<br>development | 6 | 0.0115 |
| 40 | GO:0016241 | regulation of<br>macroautophagy | 6 | 0.0123 |
| 41 | GO:0043200 | response to amino acid | 4 | 0.0125 |
| 42 | GO:0051897 | positive regulation of<br>phosphatidylinositol<br>3-kinase/protein kinase B<br>signal transduction | 12 | 0.0131 |
| 43 | GO:0034440 | lipid oxidation | 3 | 0.0140 |
| 44 | GO:0060964 | regulation of<br>miRNA-mediated gene<br>silencing | 3 | 0.0140 |
| 45 | GO:0002090 | regulation of receptor<br>internalization | 3 | 0.0140 |
| 46 | GO:0030311 | poly-N-acetyllactosamine<br>biosynthetic process | 3 | 0.0140 |
| 47 | GO:1902412 | regulation of mitotic<br>cytokinesis | 3 | 0.0140 |
| 48 | GO:0051924 | regulation of calcium ion<br>transport | 4 | 0.0142 |
| 49 | GO:0061001 | regulation of dendritic spine<br>morphogenesis | 4 | 0.0142 |
| 50 | GO:0045590 | negative regulation of<br>regulatory T cell<br>differentiation | 4 | 0.0142 |
| 51 | GO:0001525 | angiogenesis | 14 | 0.0148 |
| 52 | GO:0046777 | protein autophosphorylation | 8 | 0.0151 |
| 53 | GO:0006953 | acute-phase response | 5 | 0.0153 |
| 54 | GO:0030836 | positive regulation of actin<br>filament depolymerization | 3 | 0.0177 |
| 55 | GO:0160063 | multi-pass transmembrane<br>protein insertion into ER<br>membrane | 3 | 0.0177 |
| 56 | GO:0034599 | cellular response to oxidative<br>stress | 8 | 0.0188 |
| 57 | GO:0030168 | platelet activation | 6 | 0.0193 |
| 58 | GO:0034614 | cellular response to reactive<br>oxygen species | 5 | 0.0194 |
| 59 | GO:0034314 | Arp2/3 complex-mediated<br>actin nucleation | 4 | 0.0201 |
| 60 | GO:0006511 | ubiquitin-dependent protein<br>catabolic process | 13 | 0.0202 |
| 61 | GO:0030216 | keratinocyte differentiation | 6 | 0.0217 |
| 62 | GO:0050805 | negative regulation of<br>synaptic transmission | 3 | 0.0218 |
| 63 | GO:0050804 | modulation of chemical<br>synaptic transmission | 8 | 0.0231 |
| 64 | GO:0002181 | cytoplasmic translation | 7 | 0.0255 |
| 65 | GO:0001755 | neural crest cell migration | 5 | 0.0259 |
| 66 | GO:0071526 | semaphorin-plexin signaling<br>pathway | 5 | 0.0259 |
| 67 | GO:0034341 | response to type II interferon | 5 | 0.0259 |
| 68 | GO:0051051 | negative regulation of<br>transport | 3 | 0.0263 |
| 69 | GO:0099159 | regulation of modification of postsynaptic structure | 3 | 0.0263 |
| 70 | GO:0032814 | regulation of natural killer cell<br>activation | 3 | 0.0263 |
| 71 | GO:0034329 | cell junction assembly | 3 | 0.0263 |
| 72 | GO:0003140 | determination of left/right<br>asymmetry in lateral<br>mesoderm | 3 | 0.0263 |
| 73 | GO:0021680 | cerebellar Purkinje cell layer<br>development | 3 | 0.0263 |
| 74 | GO:0032743 | positive regulation of<br>interleukin-2 production | 5 | 0.0295 |
| 75 | GO:0099550 | trans-synaptic signaling,<br>modulating synaptic<br>transmission | 3 | 0.0311 |
| 76 | GO:0010592 | positive regulation of<br>lamellipodium assembly | 4 | 0.0326 |
| 77 | GO:0043434 | response to peptide hormone | 5 | 0.0335 |
| 78 | GO:0070914 | UV-damage excision repair | 3 | 0.0361 |
| 79 | GO:1904178 | negative regulation of<br>adipose tissue development | 3 | 0.0361 |
| 80 | GO:0035493 | SNARE complex assembly | 3 | 0.0361 |
| 81 | GO:0015031 | protein transport | 21 | 0.0391 |
| 82 | GO:0034446 | substrate<br>adhesion-dependent cell<br>spreading | 5 | 0.0400 |
| 83 | GO:0045861 | negative regulation of<br>proteolysis | 4 | 0.0417 |
| 84 | GO:0043001 | Golgi to plasma membrane<br>protein transport | 4 | 0.0417 |
| 85 | GO:0061136 | regulation of proteasomal<br>protein catabolic process | 5 | 0.0423 |
| 86 | GO:0007157 | heterophilic cell-cell<br>adhesion | 5 | 0.0423 |
| 87 | GO:0030834 | regulation of actin filament<br>depolymerization | 2 | 0.0464 |
| 88 | GO:0033206 | meiotic cytokinesis | 2 | 0.0464 |
| 89 | GO:0061097 | regulation of protein tyrosine<br>kinase activity | 2 | 0.0464 |
| 90 | GO:0046642 | negative regulation of<br>alpha-beta T cell proliferation | 2 | 0.0464 |
| 91 | GO:0015723 | bilirubin transport | 2 | 0.0464 |
| 92 | GO:1990386 | mitotic cleavage furrow<br>ingression | 2 | 0.0464 |
| 93 | GO:0003065 | positive regulation of heart<br>rate by epinephrine | 2 | 0.0464 |
| 94 | GO:0016344 | meiotic chromosome<br>movement towards spindle<br>pole | 2 | 0.0464 |
| 95 | GO:0042167 | heme catabolic process | 3 | 0.0472 |
| Note: A total of 95 significantly enriched GO BP terms (P |  |  |  |  |
| < 0.05).<br>Enrichment<br>analysis was<br>performed<br>using DAVID<br>6.8, with<br>UniProt<br>Accession<br>as the<br>identifier<br>and Homo<br>sapiens<br>whole<br>genome as<br>the<br>background. |  |  |  |  |

Subject R16 identified a total of 38 significantly enriched GO BP terms at 22–24 weeks of gestation (Table 7), the fewest among the three at this time point. The organ/tissue development-related terms included: angiogenesis (P = 1.80×10⁻³), cardiac muscle cell proliferation (P = 3.09×10⁻²), and axon guidance (P = 4.84×10⁻²).

**Table 7.**
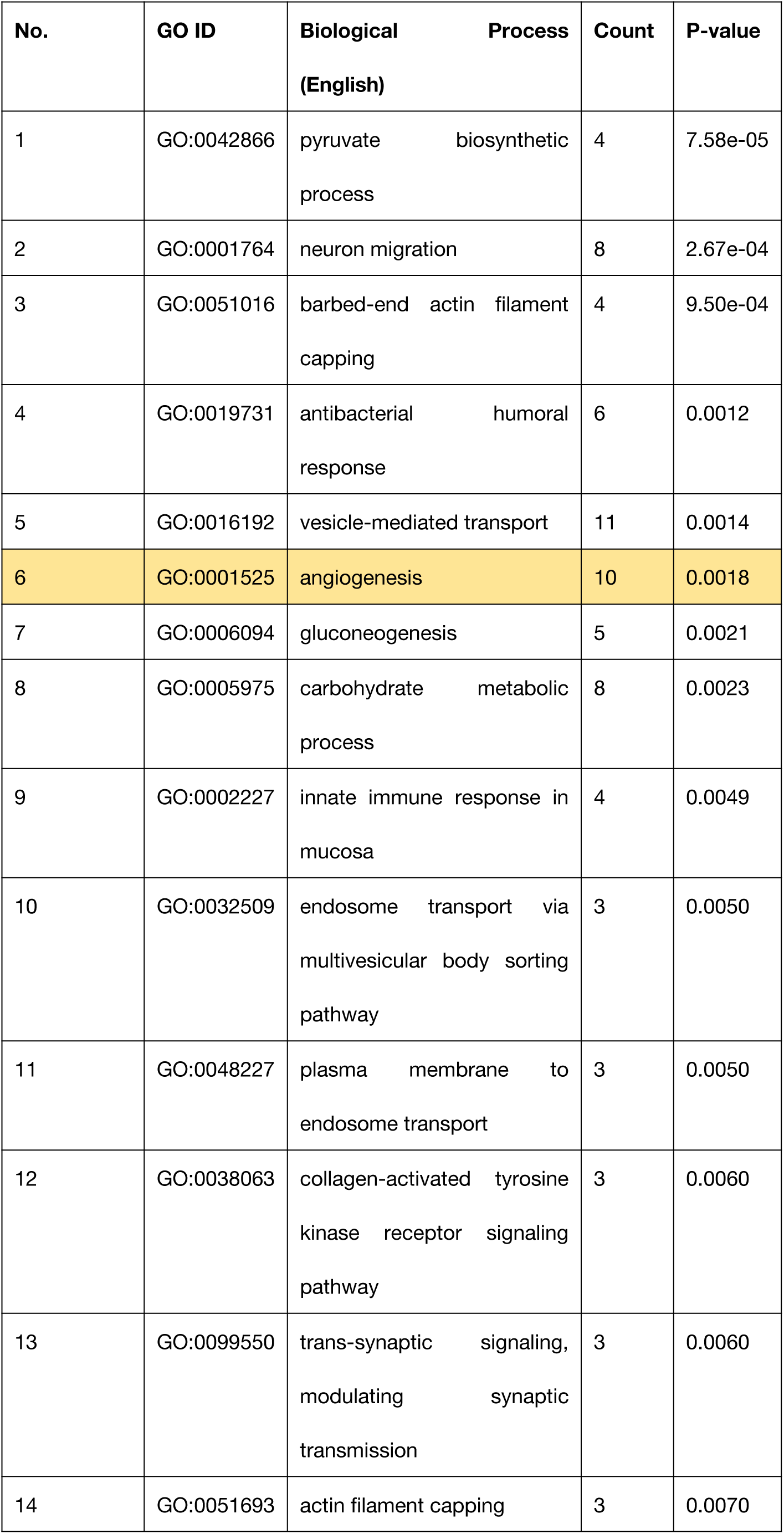
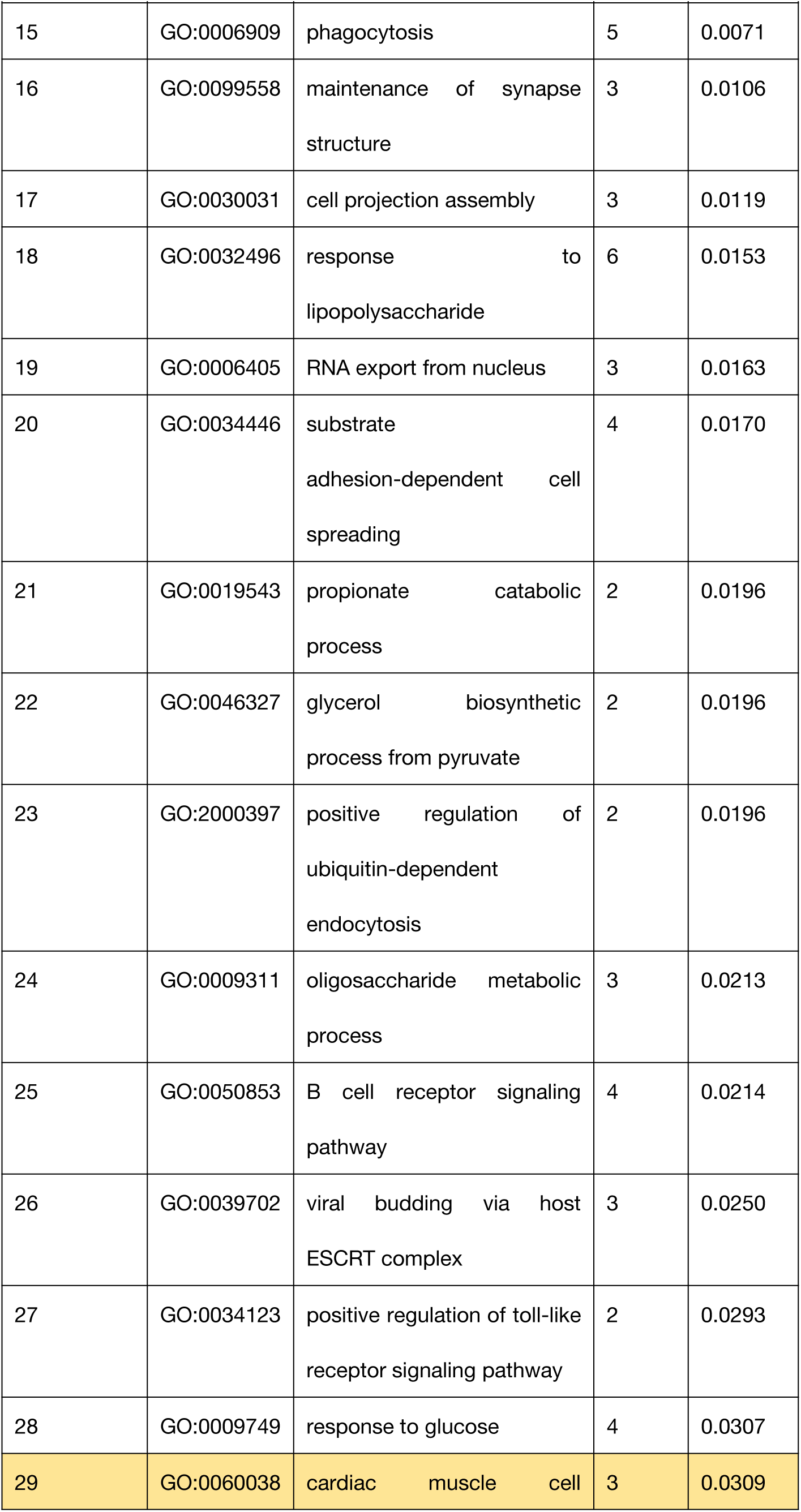
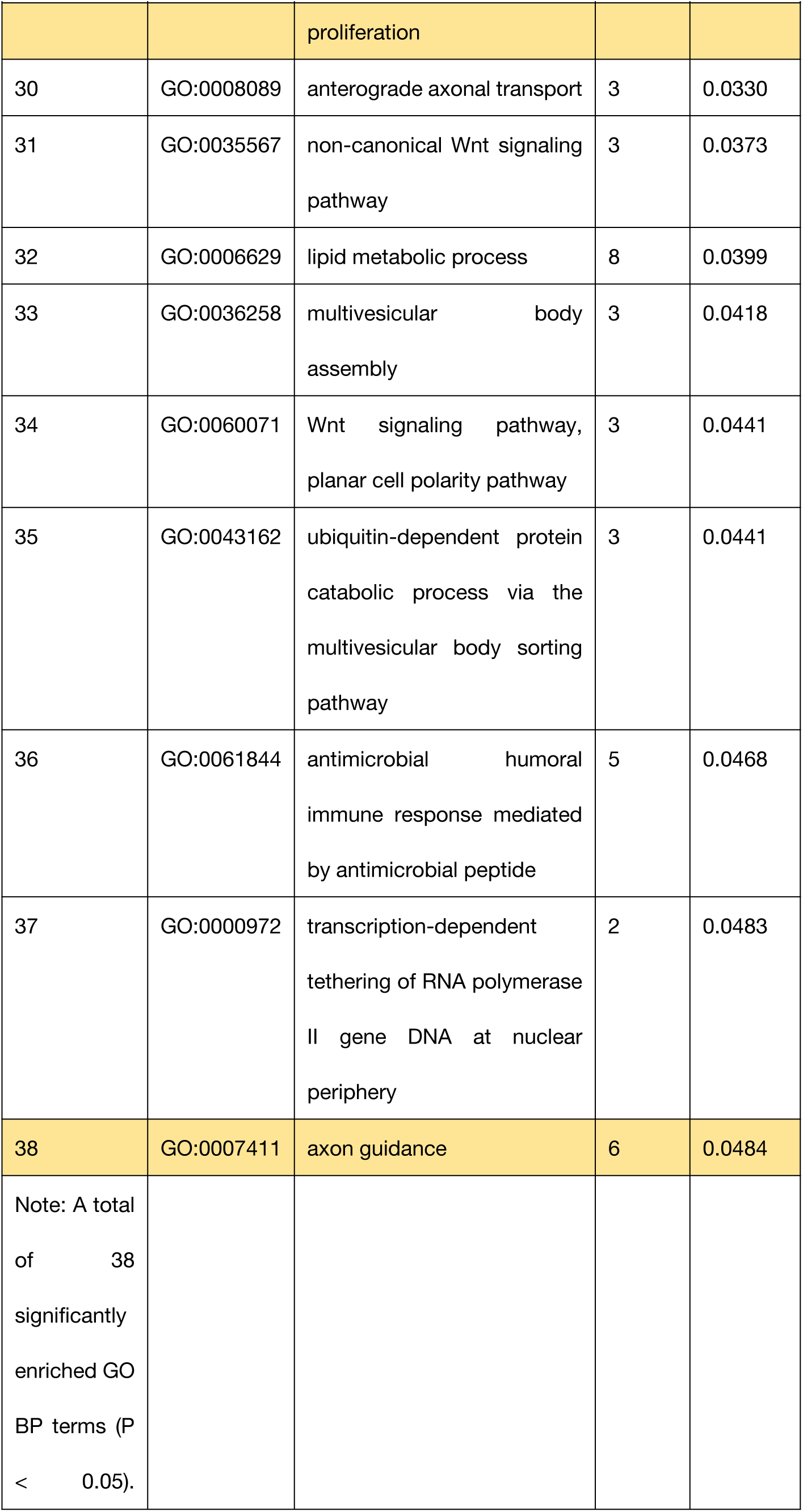
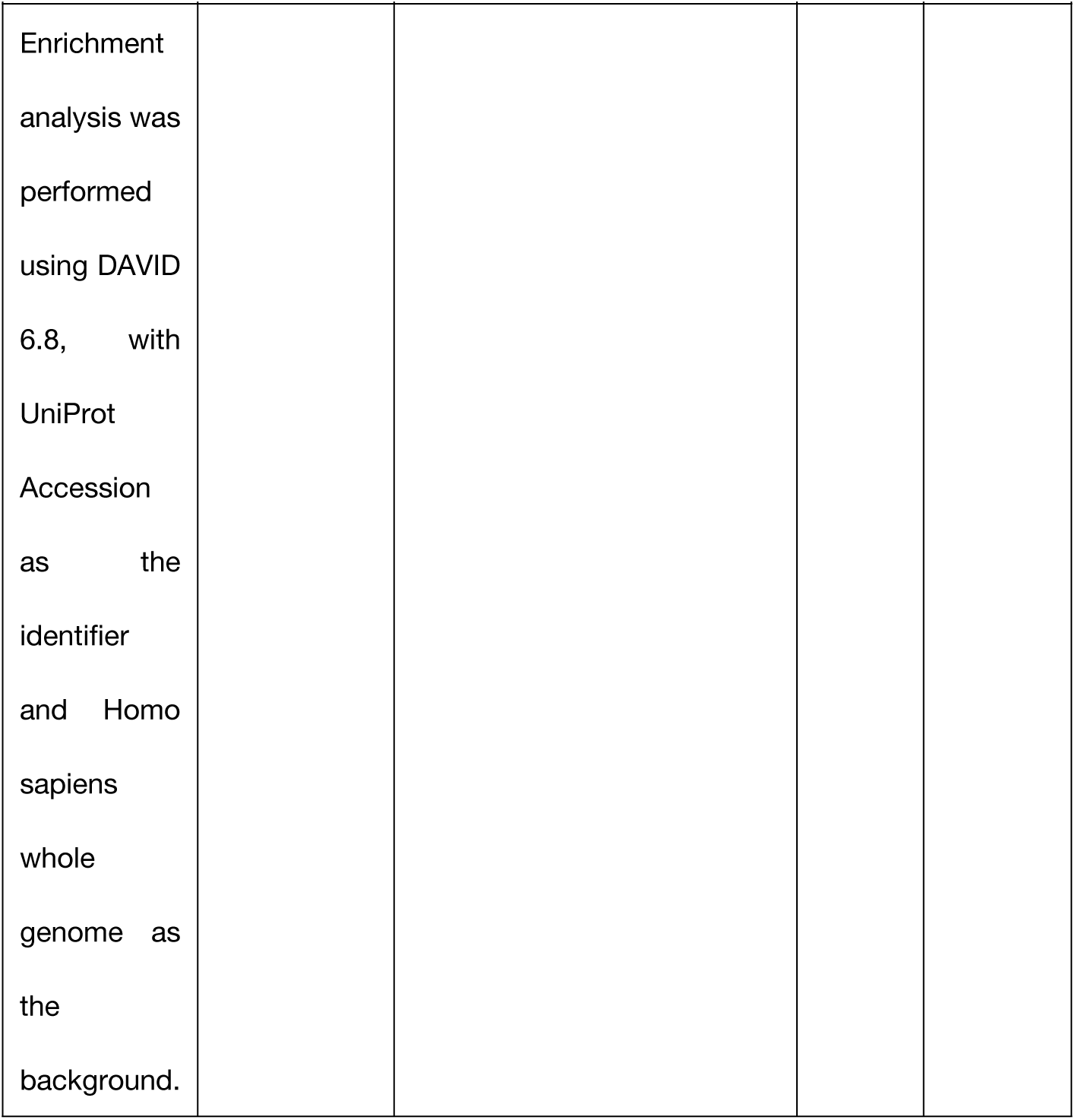
GO BP enrichment terms for R16 at 22–24 weeks of gestation (P < 0.05, 38 terms in total).

| No. | GO ID | Biological Process<br>(English) | Count | P-value |
| --- | --- | --- | --- | --- |
| 1 | GO:0042866 | pyruvate biosynthetic process | 4 | 7.58e-05 |
| 2 | GO:0001764 | neuron migration | 8 | 2.67e-04 |
| 3 | GO:0051016 | barbed-end actin filament capping | 4 | 9.50e-04 |
| 4 | GO:0019731 | antibacterial humoral response | 6 | 0.0012 |
| 5 | GO:0016192 | vesicle-mediated transport | 11 | 0.0014 |
| 6 | GO:0001525 | angiogenesis | 10 | 0.0018 |
| 7 | GO:0006094 | gluconeogenesis | 5 | 0.0021 |
| 8 | GO:0005975 | carbohydrate metabolic process | 8 | 0.0023 |
| 9 | GO:0002227 | innate immune response in mucosa | 4 | 0.0049 |
| 10 | GO:0032509 | endosome transport via multivesicular body sorting pathway | 3 | 0.0050 |
| 11 | GO:0048227 | plasma membrane to endosome transport | 3 | 0.0050 |
| 12 | GO:0038063 | collagen-activated tyrosine kinase receptor signaling pathway | 3 | 0.0060 |
| 13 | GO:0099550 | trans-synaptic signaling, modulating synaptic transmission | 3 | 0.0060 |
| 14 | GO:0051693 | actin filament capping | 3 | 0.0070 |
| 15 | GO:0006909 | phagocytosis | 5 | 0.0071 |
| 16 | GO:0099558 | maintenance of synapse<br>structure | 3 | 0.0106 |
| 17 | GO:0030031 | cell projection assembly | 3 | 0.0119 |
| 18 | GO:0032496 | response to<br>lipopolysaccharide | 6 | 0.0153 |
| 19 | GO:0006405 | RNA export from nucleus | 3 | 0.0163 |
| 20 | GO:0034446 | substrate<br>adhesion-dependent cell<br>spreading | 4 | 0.0170 |
| 21 | GO:0019543 | propionate catabolic<br>process | 2 | 0.0196 |
| 22 | GO:0046327 | glycerol biosynthetic<br>process from pyruvate | 2 | 0.0196 |
| 23 | GO:2000397 | positive regulation of<br>ubiquitin-dependent<br>endocytosis | 2 | 0.0196 |
| 24 | GO:0009311 | oligosaccharide metabolic<br>process | 3 | 0.0213 |
| 25 | GO:0050853 | B cell receptor signaling<br>pathway | 4 | 0.0214 |
| 26 | GO:0039702 | viral budding via host<br>ESCRT complex | 3 | 0.0250 |
| 27 | GO:0034123 | positive regulation of toll-like<br>receptor signaling pathway | 2 | 0.0293 |
| 28 | GO:0009749 | response to glucose | 4 | 0.0307 |
| 29 | GO:0060038 | cardiac muscle cell proliferation | 3 | 0.0309 |
| 30 | GO:0008089 | anterograde axonal transport | 3 | 0.0330 |
| 31 | GO:0035567 | non-canonical Wnt signaling pathway | 3 | 0.0373 |
| 32 | GO:0006629 | lipid metabolic process | 8 | 0.0399 |
| 33 | GO:0036258 | multivesicular body assembly | 3 | 0.0418 |
| 34 | GO:0060071 | Wnt signaling pathway, planar cell polarity pathway | 3 | 0.0441 |
| 35 | GO:0043162 | ubiquitin-dependent protein catabolic process via the multivesicular body sorting pathway | 3 | 0.0441 |
| 36 | GO:0061844 | antimicrobial humoral immune response mediated by antimicrobial peptide | 5 | 0.0468 |
| 37 | GO:0000972 | transcription-dependent tethering of RNA polymerase II gene DNA at nuclear periphery | 2 | 0.0483 |
| 38 | GO:0007411 | axon guidance | 6 | 0.0484 |
| Note: A total of 38 significantly enriched GO BP terms (P < 0.05). |  |  |  |  |
| Enrichment analysis was performed using DAVID 6.8, with UniProt Accession as the identifier and Homo sapiens whole genome as the background. |  |  |  |  |

Subject R6 identified only 28 significantly enriched GO BP terms at 32–34 weeks of gestation (Table 8), a substantial decrease compared with the 22–24 weeks time point (269 terms), suggesting that this individual’s proteomic signals were markedly weakened at the 32–34 weeks time point. Only one organ/tissue development-related term was identified—osteoclast differentiation (P = 3.88×10⁻²)—indicating that R6’s organ-development-related signals had largely subsided by the 32–34 weeks time point. The remaining terms were mainly cell adhesion and extracellular matrix remodeling (the most significant being cell–cell adhesion, P = 4.90×10⁻⁴).

**Table 8.** GO BP enrichment terms for R6 at 32–34 weeks of gestation (P < 0.05, 28 terms in total).

| No. | GO ID | Biological Process<br>(English) | Count | P-value |
| --- | --- | --- | --- | --- |
| 1 | GO:0098609 | cell-cell adhesion | 8 | 4.90e-04 |
| 2 | GO:0051289 | protein homotetramerization | 5 | 7.94e-04 |
| 3 | GO:0034332 | adherens junction<br>organization | 4 | 0.0011 |
| 4 | GO:0051180 | vitamin transport | 3 | 0.0016 |
| 5 | GO:0007043 | cell-cell junction assembly | 4 | 0.0017 |
| 6 | GO:0044331 | cell-cell adhesion mediated<br>by cadherin | 4 | 0.0019 |
| 7 | GO:0006958 | complement activation,<br>classical pathway | 4 | 0.0023 |
| 8 | GO:0016339 | calcium-dependent cell-cell<br>adhesion | 4 | 0.0029 |
| 9 | GO:0006979 | response to oxidative stress | 6 | 0.0045 |
| 10 | GO:0000902 | cell morphogenesis | 5 | 0.0049 |
| 11 | GO:0007156 | homophilic cell-cell<br>adhesion | 6 | 0.0050 |
| 12 | GO:0042157 | lipoprotein metabolic<br>process | 3 | 0.0086 |
| 13 | GO:0019731 | antibacterial humoral<br>response | 4 | 0.0129 |
| 14 | GO:0008203 | cholesterol metabolic<br>process | 4 | 0.0162 |
| 15 | GO:0006956 | complement activation | 3 | 0.0172 |
| 16 | GO:0032375 | negative regulation of<br>cholesterol transport | 2 | 0.0242 |
| 17 | GO:0008104 | intracellular protein<br>localization | 5 | 0.0245 |
| 18 | GO:0043152 | induction of bacterial<br>agglutination | 2 | 0.0301 |
| 19 | GO:0007166 | cell surface receptor<br>signaling pathway | 7 | 0.0328 |
| 20 | GO:0007155 | cell adhesion | 9 | 0.0339 |
| 21 | GO:0006084 | acetyl-CoA metabolic<br>process | 2 | 0.0361 |
| 22 | GO:0070830 | bicellular tight junction<br>assembly | 3 | 0.0388 |
| 23 | GO:0030316 | osteoclast differentiation | 3 | 0.0388 |
| 24 | GO:0002719 | negative regulation of<br>cytokine production<br>involved in immune<br>response | 2 | 0.0420 |
| 25 | GO:0007080 | mitotic metaphase<br>chromosome alignment | 3 | 0.0445 |
| 26 | GO:0032760 | positive regulation of tumor<br>necrosis factor production | 4 | 0.0454 |
| 27 | GO:0006629 | lipid metabolic process | 6 | 0.0470 |
| 28 | GO:0046949 | fatty-acyl-CoA biosynthetic<br>process | 2 | 0.0478 |
| Note: A total of 28 significantly enriched GO BP terms ($P < 0.05$). Enrichment analysis was |  |  |  |  |
| performed using DAVID 6.8, with UniProt Accession as the identifier and Homo sapiens whole genome as the background. |  |  |  |  |

Subject R15 identified a total of 96 significantly enriched GO BP terms at 32–34 weeks of gestation (Table 9), roughly the same number as at the 22–24 weeks time point (95 terms). A total of 19 organ/tissue development or morphogenesis-related terms were identified, including: axon guidance (P = 2.62×10⁻⁵), heart development (P = 3.04×10⁻³), hair follicle development (P = 4.24×10⁻³), osteoblast differentiation (P = 6.25×10⁻³), endothelial cell differentiation (P = 8.46×10⁻³), neuron projection development (P = 8.89×10⁻³), endodermal cell differentiation (P = 9.37×10⁻³), retinal ganglion cell axon guidance (P = 1.31×10⁻²), synapse assembly (P = 1.37×10⁻²), angiogenesis (P = 1.66×10⁻²), nervous system development (P = 1.75×10⁻²), erythrocyte differentiation (P = 2.19×10⁻²), substantia nigra development (P = 2.56×10⁻²), epithelial cell differentiation (P = 3.27×10⁻²) and epidermis development (P = 3.27×10⁻²), kidney development (P = 3.29×10⁻²), and aorta development (P = 3.71×10⁻²), overall showing a persistent broad-peak developmental signal pattern.

**Table 9.** GO BP enrichment terms for R15 at 32–34 weeks of gestation (P < 0.05, 96 terms in total).

| No. | GO ID | Biological Process<br>(English) | Count | P-value |
| --- | --- | --- | --- | --- |
| 1 | GO:0002227 | innate immune response in mucosa | 15 | 2.37e-14 |
| 2 | GO:0031507 | heterochromatin formation | 23 | 1.13e-12 |
| 3 | GO:0006325 | chromatin organization | 32 | 2.79e-10 |
| 4 | GO:0019731 | antibacterial humoral<br>response | 16 | 6.01e-10 |
| 5 | GO:0098609 | cell-cell adhesion | 25 | 1.20e-09 |
| 6 | GO:0061844 | antimicrobial humoral<br>immune response mediated<br>by antimicrobial peptide | 18 | 3.03e-08 |
| 7 | GO:0007155 | cell adhesion | 39 | 8.53e-08 |
| 8 | GO:0007156 | homophilic cell-cell<br>adhesion | 19 | 3.35e-07 |
| 9 | GO:0050830 | defense response to<br>Gram-positive bacterium | 17 | 3.62e-07 |
| 10 | GO:0016197 | endosomal transport | 13 | 1.05e-06 |
| 11 | GO:0015031 | protein transport | 33 | 4.40e-06 |
| 12 | GO:0034315 | regulation of Arp2/3<br>complex-mediated actin<br>nucleation | 7 | 1.29e-05 |
| 13 | GO:0007411 | axon guidance | 17 | 2.62e-05 |
| 14 | GO:0006334 | nucleosome assembly | 15 | 3.57e-05 |
| 15 | GO:0030335 | positive regulation of cell<br>migration | 20 | 9.46e-05 |
| 16 | GO:0032456 | endocytic recycling | 10 | 2.75e-04 |
| 17 | GO:0050919 | negative chemotaxis | 7 | 3.47e-04 |
| 18 | GO:0042104 | positive regulation of<br>activated T cell proliferation | 6 | 4.75e-04 |
| 19 | GO:0030036 | actin cytoskeleton organization | 16 | 6.59e-04 |
| 20 | GO:0006914 | autophagy | 14 | 6.70e-04 |
| 21 | GO:0051897 | positive regulation of phosphatidylinositol 3-kinase/protein kinase B signal transduction | 15 | 6.90e-04 |
| 22 | GO:0051489 | regulation of filopodium assembly | 5 | 7.54e-04 |
| 23 | GO:0007165 | signal transduction | 53 | 8.65e-04 |
| 24 | GO:0042147 | retrograde transport, endosome to Golgi | 9 | 0.0015 |
| 25 | GO:0007160 | cell-matrix adhesion | 10 | 0.0017 |
| 26 | GO:0007159 | leukocyte cell-cell adhesion | 6 | 0.0018 |
| 27 | GO:0030199 | collagen fibril organization | 8 | 0.0021 |
| 28 | GO:2001237 | negative regulation of extrinsic apoptotic signaling pathway | 6 | 0.0022 |
| 29 | GO:0044331 | cell-cell adhesion mediated by cadherin | 6 | 0.0025 |
| 30 | GO:0034314 | Arp2/3 complex-mediated actin nucleation | 5 | 0.0027 |
| 31 | GO:0007507 | heart development | 14 | 0.0030 |
| 32 | GO:0001942 | hair follicle development | 6 | 0.0042 |
| 33 | GO:0043542 | endothelial cell migration | 6 | 0.0047 |
| 34 | GO:0030866 | cortical actin cytoskeleton organization | 5 | 0.0048 |
| 35 | GO:0002250 | adaptive immune response | 23 | 0.0050 |
| 36 | GO:0016339 | calcium-dependent cell-cell<br>adhesion | 6 | 0.0051 |
| 37 | GO:0031032 | actomyosin structure<br>organization | 5 | 0.0054 |
| 38 | GO:0001649 | osteoblast differentiation | 10 | 0.0063 |
| 39 | GO:0001938 | positive regulation of<br>endothelial cell proliferation | 7 | 0.0070 |
| 40 | GO:0045446 | endothelial cell<br>differentiation | 4 | 0.0085 |
| 41 | GO:0043410 | positive regulation of MAPK<br>cascade | 13 | 0.0086 |
| 42 | GO:0031175 | neuron projection<br>development | 10 | 0.0089 |
| 43 | GO:0008284 | positive regulation of cell<br>population proliferation | 23 | 0.0093 |
| 44 | GO:0035987 | endodermal cell<br>differentiation | 5 | 0.0094 |
| 45 | GO:0006887 | exocytosis | 9 | 0.0108 |
| 46 | GO:0090263 | positive regulation of<br>canonical Wnt signaling<br>pathway | 9 | 0.0113 |
| 47 | GO:0061436 | establishment of skin barrier | 5 | 0.0114 |
| 48 | GO:0007413 | axonal fasciculation | 4 | 0.0114 |
| 49 | GO:0031290 | retinal ganglion cell axon<br>guidance | 4 | 0.0131 |
| 50 | GO:0007416 | synapse assembly | 7 | 0.0137 |
| 51 | GO:0090148 | membrane fission | 5 | 0.0162 |
| 52 | GO:0001525 | angiogenesis | 14 | 0.0166 |
| 53 | GO:0007399 | nervous system<br>development | 22 | 0.0175 |
| 54 | GO:0008354 | germ cell migration | 3 | 0.0183 |
| 55 | GO:0031295 | T cell costimulation | 5 | 0.0206 |
| 56 | GO:0032091 | negative regulation of<br>protein binding | 4 | 0.0211 |
| 57 | GO:1900273 | positive regulation of<br>long-term synaptic<br>potentiation | 4 | 0.0211 |
| 58 | GO:0030307 | positive regulation of cell<br>growth | 7 | 0.0218 |
| 59 | GO:0000281 | mitotic cytokinesis | 6 | 0.0219 |
| 60 | GO:0046718 | symbiont entry into host cell | 6 | 0.0219 |
| 61 | GO:0030218 | erythrocyte differentiation | 6 | 0.0219 |
| 62 | GO:0035425 | autocrine signaling | 3 | 0.0226 |
| 63 | GO:0002544 | chronic inflammatory<br>response | 3 | 0.0226 |
| 64 | GO:0050821 | protein stabilization | 13 | 0.0228 |
| 65 | GO:0032922 | circadian regulation of gene<br>expression | 6 | 0.0232 |
| 66 | GO:0010824 | regulation of centrosome<br>duplication | 4 | 0.0234 |
| 67 | GO:0001818 | negative regulation of<br>cytokine production | 5 | 0.0238 |
| 68 | GO:0021762 | substantia nigra<br>development | 5 | 0.0256 |
| 69 | GO:0051469 | vesicle fusion with vacuole | 3 | 0.0272 |
| 70 | GO:0036010 | protein localization to endosome | 3 | 0.0272 |
| 71 | GO:0070527 | platelet aggregation | 5 | 0.0274 |
| 72 | GO:1901673 | regulation of mitotic spindle assembly | 4 | 0.0285 |
| 73 | GO:0009117 | nucleotide metabolic process | 4 | 0.0285 |
| 74 | GO:0006796 | phosphate-containing compound metabolic process | 4 | 0.0285 |
| 75 | GO:0061763 | multivesicular body-lysosome fusion | 3 | 0.0321 |
| 76 | GO:0030855 | epithelial cell differentiation | 7 | 0.0327 |
| 77 | GO:0008544 | epidermis development | 7 | 0.0327 |
| 78 | GO:0001822 | kidney development | 8 | 0.0329 |
| 79 | GO:0006096 | glycolytic process | 5 | 0.0333 |
| 80 | GO:0051491 | positive regulation of filopodium assembly | 4 | 0.0371 |
| 81 | GO:0035904 | aorta development | 4 | 0.0371 |
| 82 | GO:0001837 | epithelial to mesenchymal transition | 5 | 0.0376 |
| 83 | GO:0006955 | immune response | 22 | 0.0384 |
| 84 | GO:0006749 | glutathione metabolic process | 5 | 0.0398 |
| 85 | GO:0006893 | Golgi to plasma membrane transport | 4 | 0.0403 |
| 86 | GO:0051493 | regulation of cytoskeleton organization | 4 | 0.0403 |
| 87 | GO:0034113 | heterotypic cell-cell adhesion | 4 | 0.0403 |
| 88 | GO:0071679 | commissural neuron axon guidance | 3 | 0.0429 |
| 89 | GO:0090303 | positive regulation of wound healing | 4 | 0.0435 |
| 90 | GO:0034332 | adherens junction organization | 4 | 0.0435 |
| 91 | GO:0036258 | multivesicular body assembly | 4 | 0.0435 |
| 92 | GO:0007157 | heterophilic cell-cell adhesion | 5 | 0.0446 |
| 93 | GO:0009611 | response to wounding | 6 | 0.0465 |
| 94 | GO:0035924 | cellular response to vascular endothelial growth factor stimulus | 4 | 0.0469 |
| 95 | GO:2000536 | negative regulation of entry of bacterium into host cell | 2 | 0.0472 |
| 96 | GO:0070488 | neutrophil aggregation | 2 | 0.0472 |
| Note: A total of 96 significantly enriched GO BP terms (P |  |  |  |  |
| < 0.05).<br>Enrichment<br>analysis was<br>performed<br>using DAVID<br>6.8, with<br>UniProt<br>Accession<br>as the<br>identifier<br>and Homo<br>sapiens<br>whole<br>genome as<br>the<br>background. |  |  |  |  |

Subject R16 identified a total of 140 significantly enriched GO BP terms at 32–34 weeks of gestation (Table 10), the largest number among the three pregnant women at this time point, in sharp contrast to the weaker signal pattern at the 22–24 weeks time point (38 terms). A total of 18 organ/tissue development or morphogenesis-related terms were identified, with neural and vascular developmental signals being the most prominent: angiogenesis (P = 3.64×10⁻³), epithelial cell differentiation (P = 3.89×10⁻³), neuron projection development (P = 8.50×10⁻³), commissural neuron axon guidance (P = 1.35×10⁻²), nervous system development (P = 1.48×10⁻²), axon guidance (P = 1.69×10⁻²), in utero embryonic development (P = 1.70×10⁻²), venous blood vessel morphogenesis (P = 2.53×10⁻²), head morphogenesis (P = 4.14×10⁻²), and digestive tract development (P = 3.57×10⁻²), suggesting that R16’s main developmental signal peak was at 32–34 weeks of gestation, showing a delayed-peak pattern.

**Table 10.** GO BP enrichment terms for R16 at 32–34 weeks of gestation (P < 0.05, 140 terms in total).

| No. | GO ID | Biological Process<br>(English) | Count | P-value |
| --- | --- | --- | --- | --- |
| 1 | GO:0007155 | cell adhesion | 54 | 8.44e-09 |
| 2 | GO:0030335 | positive regulation of cell migration | 32 | 1.82e-07 |
| 3 | GO:0007156 | homophilic cell-cell adhesion | 24 | 2.27e-07 |
| 4 | GO:0006749 | glutathione metabolic process | 13 | 4.91e-07 |
| 5 | GO:0098609 | cell-cell adhesion | 25 | 3.95e-06 |
| 6 | GO:0007165 | signal transduction | 83 | 1.66e-05 |
| 7 | GO:0015031 | protein transport | 42 | 2.20e-05 |
| 8 | GO:0006887 | exocytosis | 16 | 6.52e-05 |
| 9 | GO:0031623 | receptor internalization | 10 | 1.09e-04 |
| 10 | GO:0045087 | innate immune response | 40 | 1.65e-04 |
| 11 | GO:0043410 | positive regulation of MAPK cascade | 21 | 3.09e-04 |
| 12 | GO:0006955 | immune response | 39 | 4.72e-04 |
| 13 | GO:0006898 | receptor-mediated endocytosis | 11 | 8.70e-04 |
| 14 | GO:0016477 | cell migration | 23 | 8.79e-04 |
| 15 | GO:0007157 | heterophilic cell-cell adhesion | 9 | 9.94e-04 |
| 16 | GO:0006893 | Golgi to plasma membrane transport | 7 | 9.97e-04 |
| 17 | GO:0043066 | negative regulation of apoptotic process | 36 | 0.0011 |
| 18 | GO:0016197 | endosomal transport | 11 | 0.0014 |
| 19 | GO:0042177 | negative regulation of protein catabolic process | 8 | 0.0014 |
| 20 | GO:0031179 | peptide modification | 4 | 0.0016 |
| 21 | GO:0008286 | insulin receptor signaling pathway | 10 | 0.0016 |
| 22 | GO:0007596 | blood coagulation | 12 | 0.0016 |
| 23 | GO:0032869 | cellular response to insulin stimulus | 12 | 0.0016 |
| 24 | GO:0032456 | endocytic recycling | 11 | 0.0016 |
| 25 | GO:0019221 | cytokine-mediated signaling pathway | 15 | 0.0016 |
| 26 | GO:1902430 | negative regulation of amyloid-beta formation | 7 | 0.0019 |
| 27 | GO:0007169 | cell surface receptor protein tyrosine kinase signaling pathway | 13 | 0.0022 |
| 28 | GO:0050819 | negative regulation of coagulation | 4 | 0.0024 |
| 29 | GO:0006635 | fatty acid beta-oxidation | 8 | 0.0026 |
| 30 | GO:0008284 | positive regulation of cell population proliferation | 34 | 0.0027 |
| 31 | GO:0010544 | negative regulation of platelet activation | 4 | 0.0036 |
| 32 | GO:0001525 | angiogenesis | 21 | 0.0036 |
| 33 | GO:0001523 | retinoid metabolic process | 7 | 0.0037 |
| 34 | GO:0030855 | epithelial cell differentiation | 11 | 0.0039 |
| 35 | GO:0002250 | adaptive immune response | 32 | 0.0040 |
| 36 | GO:0048146 | positive regulation of fibroblast proliferation | 8 | 0.0040 |
| 37 | GO:2000630 | positive regulation of miRNA metabolic process | 3 | 0.0040 |
| 38 | GO:0030334 | regulation of cell migration | 12 | 0.0043 |
| 39 | GO:0009725 | response to hormone | 8 | 0.0044 |
| 40 | GO:0042572 | retinol metabolic process | 8 | 0.0044 |
| 41 | GO:0008037 | cell recognition | 4 | 0.0050 |
| 42 | GO:0046628 | positive regulation of insulin receptor signaling pathway | 5 | 0.0056 |
| 43 | GO:0006691 | leukotriene metabolic process | 5 | 0.0056 |
| 44 | GO:0032956 | regulation of actin cytoskeleton organization | 11 | 0.0063 |
| 45 | GO:0051897 | positive regulation of phosphatidylinositol 3-kinase/protein kinase B signal transduction | 17 | 0.0064 |
| 46 | GO:0042147 | retrograde transport, endosome to Golgi | 10 | 0.0065 |
| 47 | GO:0090331 | negative regulation of platelet aggregation | 4 | 0.0066 |
| 48 | GO:0009395 | phospholipid catabolic process | 5 | 0.0067 |
| 49 | GO:0043087 | regulation of GTPase activity | 6 | 0.0069 |
| 50 | GO:0016339 | calcium-dependent cell-cell adhesion | 7 | 0.0074 |
| 51 | GO:0071267 | L-methionine salvage | 3 | 0.0078 |
| 52 | GO:0042060 | wound healing | 9 | 0.0081 |
| 53 | GO:0070527 | platelet aggregation | 7 | 0.0082 |
| 54 | GO:0031175 | neuron projection development | 13 | 0.0085 |
| 55 | GO:0043407 | negative regulation of MAP kinase activity | 4 | 0.0086 |
| 56 | GO:0046626 | regulation of insulin receptor signaling pathway | 4 | 0.0086 |
| 57 | GO:0030155 | regulation of cell adhesion | 8 | 0.0092 |
| 58 | GO:0032006 | regulation of TOR signaling | 5 | 0.0094 |
| 59 | GO:0034315 | regulation of Arp2/3 complex-mediated actin nucleation | 5 | 0.0094 |
| 60 | GO:0097190 | apoptotic signaling pathway | 9 | 0.0094 |
| 61 | GO:0006120 | mitochondrial electron transport, NADH to ubiquinone | 7 | 0.0100 |
| 62 | GO:0042776 | proton motive force-driven mitochondrial ATP<br>synthesis | 8 | 0.0108 |
| 63 | GO:0017015 | regulation of transforming<br>growth factor beta receptor<br>signaling pathway | 5 | 0.0109 |
| 64 | GO:0046627 | negative regulation of<br>insulin receptor signaling<br>pathway | 7 | 0.0110 |
| 65 | GO:0007159 | leukocyte cell-cell adhesion | 6 | 0.0112 |
| 66 | GO:0008217 | regulation of blood<br>pressure | 9 | 0.0116 |
| 67 | GO:0006631 | fatty acid metabolic<br>process | 12 | 0.0116 |
| 68 | GO:0036211 | protein modification<br>process | 8 | 0.0126 |
| 69 | GO:1900273 | positive regulation of<br>long-term synaptic<br>potentiation | 5 | 0.0126 |
| 70 | GO:0034314 | Arp2/3 complex-mediated<br>actin nucleation | 5 | 0.0126 |
| 71 | GO:1904504 | positive regulation of<br>lipophagy | 3 | 0.0127 |
| 72 | GO:0071679 | commissural neuron axon<br>guidance | 4 | 0.0135 |
| 73 | GO:0072344 | rescue of stalled cytosolic<br>ribosome | 6 | 0.0139 |
| 74 | GO:0071356 | cellular response to tumor necrosis factor | 10 | 0.0141 |
| 75 | GO:0007399 | nervous system development | 31 | 0.0148 |
| 76 | GO:0035556 | intracellular signal transduction | 30 | 0.0155 |
| 77 | GO:0006751 | glutathione catabolic process | 4 | 0.0164 |
| 78 | GO:0007411 | axon guidance | 15 | 0.0169 |
| 79 | GO:0001701 | in utero embryonic development | 17 | 0.0170 |
| 80 | GO:0042760 | very long-chain fatty acid catabolic process | 3 | 0.0186 |
| 81 | GO:0090166 | Golgi disassembly | 3 | 0.0186 |
| 82 | GO:0090263 | positive regulation of canonical Wnt signaling pathway | 11 | 0.0195 |
| 83 | GO:0070593 | dendrite self-avoidance | 4 | 0.0197 |
| 84 | GO:0006892 | post-Golgi vesicle-mediated transport | 4 | 0.0197 |
| 85 | GO:0043116 | negative regulation of vascular permeability | 4 | 0.0197 |
| 86 | GO:0030593 | neutrophil chemotaxis | 7 | 0.0199 |
| 87 | GO:0045197 | establishment or maintenance of epithelial cell apical/basal polarity | 5 | 0.0211 |
| 88 | GO:0010332 | response to gamma radiation | 5 | 0.0211 |
| 89 | GO:0030866 | cortical actin cytoskeleton organization | 5 | 0.0211 |
| 90 | GO:1902600 | proton transmembrane transport | 14 | 0.0219 |
| 91 | GO:0043588 | skin development | 6 | 0.0225 |
| 92 | GO:0022900 | electron transport chain | 4 | 0.0233 |
| 93 | GO:0050727 | regulation of inflammatory response | 10 | 0.0243 |
| 94 | GO:0006914 | autophagy | 14 | 0.0246 |
| 95 | GO:0048845 | venous blood vessel morphogenesis | 3 | 0.0253 |
| 96 | GO:0007160 | cell-matrix adhesion | 10 | 0.0256 |
| 97 | GO:0006997 | nucleus organization | 5 | 0.0264 |
| 98 | GO:0034599 | cellular response to oxidative stress | 10 | 0.0269 |
| 99 | GO:0098869 | cellular oxidant detoxification | 7 | 0.0286 |
| 100 | GO:0001570 | vasculogenesis | 7 | 0.0286 |
| 101 | GO:0050807 | regulation of synapse organization | 5 | 0.0293 |
| 102 | GO:0030183 | B cell differentiation | 8 | 0.0302 |
| 103 | GO:0016192 | vesicle-mediated transport | 20 | 0.0313 |
| 104 | GO:0035234 | ectopic germ cell programmed cell death | 4 | 0.0314 |
| 105 | GO:0045494 | photoreceptor cell maintenance | 6 | 0.0315 |
| 106 | GO:0002318 | myeloid progenitor cell differentiation | 3 | 0.0330 |
| 107 | GO:0001666 | response to hypoxia | 13 | 0.0356 |
| 108 | GO:0048565 | digestive tract development | 5 | 0.0357 |
| 109 | GO:0050900 | leukocyte migration | 5 | 0.0357 |
| 110 | GO:0010761 | fibroblast migration | 4 | 0.0359 |
| 111 | GO:0033138 | positive regulation of<br>peptidyl-serine<br>phosphorylation | 4 | 0.0359 |
| 112 | GO:0042552 | myelination | 7 | 0.0372 |
| 113 | GO:0043065 | positive regulation of<br>apoptotic process | 20 | 0.0383 |
| 114 | GO:0051591 | response to cAMP | 5 | 0.0392 |
| 115 | GO:0032480 | negative regulation of type I<br>interferon production | 5 | 0.0392 |
| 116 | GO:0032731 | positive regulation of<br>interleukin-1 beta<br>production | 7 | 0.0396 |
| 117 | GO:0006935 | chemotaxis | 11 | 0.0398 |
| 118 | GO:0016236 | macroautophagy | 8 | 0.0399 |
| 119 | GO:0006693 | prostaglandin metabolic<br>process | 4 | 0.0408 |
| 120 | GO:0060323 | head morphogenesis | 3 | 0.0414 |
| 121 | GO:0038134 | ERBB2-EGFR signaling<br>pathway | 3 | 0.0414 |
| 122 | GO:0044597 | daunorubicin metabolic<br>process | 3 | 0.0414 |
| 123 | GO:0001775 | cell activation | 3 | 0.0414 |
| 124 | GO:0070973 | protein localization to<br>endoplasmic reticulum exit<br>site | 3 | 0.0414 |
| 125 | GO:0032287 | peripheral nervous system<br>myelin maintenance | 3 | 0.0414 |
| 126 | GO:0010224 | response to UV-B | 3 | 0.0414 |
| 127 | GO:0000255 | allantoin metabolic process | 3 | 0.0414 |
| 128 | GO:0010642 | negative regulation of<br>platelet-derived growth<br>factor receptor signaling<br>pathway | 3 | 0.0414 |
| 129 | GO:0050830 | defense response to<br>Gram-positive bacterium | 11 | 0.0414 |
| 130 | GO:0009060 | aerobic respiration | 7 | 0.0420 |
| 131 | GO:0034097 | response to cytokine | 6 | 0.0425 |
| 132 | GO:0007010 | cytoskeleton organization | 13 | 0.0425 |
| 133 | GO:0048678 | response to axon injury | 5 | 0.0429 |
| 134 | GO:0006139 | nucleobase-containing<br>compound metabolic<br>process | 5 | 0.0429 |
| 135 | GO:0018108 | peptidyl-tyrosine<br>phosphorylation | 5 | 0.0429 |
| 136 | GO:0035329 | hippo signaling | 4 | 0.0460 |
| 137 | GO:0016485 | protein processing | 10 | 0.0462 |
| 138 | GO:0050808 | synapse organization | 7 | 0.0473 |
| 139 | GO:0071222 | cellular response to<br>lipopolysaccharide | 13 | 0.0486 |
| 140 | GO:0070374 | positive regulation of ERK1<br>and ERK2 cascade | 14 | 0.0489 |
| Note: A total of 140 significantly enriched GO BP terms (P < 0.05). Enrichment analysis was performed using DAVID 6.8, with UniProt Accession as the identifier and Homo sapiens whole genome as the background. |  |  |  |  |

Samples were independently searched and quantified in four batches using Spectronaut X. Across all four batches, a total of 5,169 unique proteins were identified (PG.Qvalue < 0.01), of which 3,673 (71.1%) were core proteins shared by all four batches. The composition of each batch and the number of identified proteins are summarized in Table 11. Given inter-batch technical variation, all differential protein screening in this study was performed by within-batch paired comparison (i.e., each pregnant woman’s pregnancy sample was compared only with the six control samples within the same batch).

**Table 11.** Overview of proteome identification across batches.

| Batch | Samples Included | Number of Identified Proteins (PG.Qvalue < 0.01) | Number of Samples within Batch |
| --- | --- | --- | --- |
| Batch1 | R6, R15, R16 (6–8 weeks of gestation) + controls C1–C6 | 3,869 | 9 |
| Batch2 | R6 (22–24 weeks + 32–34 weeks of gestation) + controls C1–C6 | 4,511 | 8 |
| Batch3 | R15, R16 (22–24 weeks of gestation) + controls C1–C6 | 4,695 | 8 |
| Batch4 | R15, R16 (32–34 weeks of gestation) + controls C1–C6 | 4,739 | 8 |
| Total (union of 4 batches) | — | 5,169 (unique proteins) | — |
| Common to all 4 batches (intersection) | — | 3,673 (71.1%) | — |

## 4. Discussion

Using a one-versus-many (one pregnant woman vs. six non-pregnant women) urinary proteomic analytical framework, this study successfully detected GO Biological Process enrichment signals at the single-sample level that are highly consistent with the physiological processes of normal pregnancy. Across all nine samples, a large number of organ, tissue development, or morphogenesis-related terms were identified, covering multiple organ systems including nervous, vascular, cardiac, skeletal, renal, pulmonary, digestive, sensory, and skin. Because there should be no continuously active organ morphogenesis in pregnant women, the widespread detection of these terms strongly suggests that the signals originate from the developing fetus—that is, the urinary proteome can capture fetal development signals. Precisely because urine need not maintain internal homeostasis, subtle changes in multiple bodily systems can be disproportionately amplified in urine, so that even at the single-sample level, the proteomic signals induced by pregnancy physiological remodeling remain clear enough to be detected by differential protein analysis.

From a biological temporal perspective, among the GO BP pathways detected at the three sampling time points, the first sampling (at a time point around 6 – 8 weeks of gestation) was dominated by maternal physiological adaptation terms (immune regulation, metabolic remodeling, cytoskeletal reorganization), yet all three pregnant women already exhibited several pathways directly related to early embryonic development. The number of these pathways was not large— likely not because the signals themselves were weak, but because of two main limiting factors: the GO database annotation for very early human embryonic development remains rather sparse, and at 6 – 8 weeks of gestation most organ systems had only just entered the initiation stage of morphogenesis, so the number of pathways was small.

The signals at the second sampling (around 22 – 24 weeks of gestation) changed markedly: all three pregnant women showed enrichment of processes including nervous system development, angiogenesis, and the development of core organs such as the heart, as well as generative or tissue cell differentiation, temporally corresponding to the critical window during which fetal organs transition from morphological differentiation to functional maturation.

The signals at the third sampling (around 32 – 34 weeks of gestation) differed among the three pregnant women: R6’s organ-development signals had mostly subsided, R15 continued to show multiple neurodevelopmental pathways, and R16’s developmental signals instead increased. Although the three showed different results, each pregnant woman independently detected fetal-development-related GO BP terms consistent with her gestational stage—a fact indicating that the urinary proteome can stably capture the biological signals of fetal development at each sampling time point, providing cross-species corroboration with the findings of Wang et al. (2025) in a rat model^[12]^; it also shows that one-versus-many analysis can avoid masking inter-individual differences under different developmental patterns and amplify the sensitivity of urinary proteomics.

The most obvious shortcoming of this study is the sample size. Urine samples from three pregnant women at nine sampling time points can demonstrate that we can see fetal development signals in urine at the individual level, but they are still insufficient for traditional population-level statistical inference. Another problem is the sparse sampling density—each pregnant woman had only three time points, with intervals of more than ten weeks, during which fetal organ development is actually a continuous process changing daily; a line drawn from only three points is coarse. If sampling density could be increased along this timeline in the future (for example, by adding a sampling point every week of gestation), there would be an opportunity to observe a truly fine-grained temporal curve of fetal development.

At a more long-term level, we believe the true value of this work lies in providing a feasible workflow for establishing a temporal atlas of the pregnancy urinary proteome. If such a temporal atlas could be systematically established, every sampling time point would have a referenceable “normal range,” and fetal developmental abnormalities deviating from the normal trajectory might then be detected as early warning signals in urine. Of course, this requires substantial work, including validation with more independent cohorts, improvements in proteomic technology (such as using organ-specific protein databases to trace signal origins), and cross-comparison with amniotic fluid or umbilical cord blood proteomes.

## 5. Conclusions

Through one-versus-many urinary proteomic comparative analysis at three pregnancy time points in three women with normal pregnancies, this study confirmed that this method can capture the temporal signals of fetal organ development at the individual level. The main findings are as follows.

First, all nine samples’ urinary proteomes independently detected GO BP terms highly relevant to fetal development at their respective stages, covering multiple organ systems including nervous, vascular, cardiac, skeletal, renal, pulmonary, digestive, sensory, and skin. The first sampling (around 6 – 8 weeks of gestation) detected fewer terms, mainly nervous system development; the second sampling (around 22 – 24 weeks of gestation) showed dense enrichment of organogenesis terms; and the third sampling (around 32 – 34 weeks of gestation) showed inter-individual differences in developmental signals, yet still enriched fetal-development-related terms. These indicate that urinary proteomics can effectively capture sensitive signals of the temporal progression of fetal development.

Second, one-versus-many analysis can detect individual-specific pathways at the single-sample level that are easily masked by group-based analysis. The fetal development timelines of the three pregnant women in this study showed marked inter-individual differences, suggesting that fetal developmental progression itself exhibits significant inter-individual variation within the physiological range. This finding has methodological implications for future pregnancy monitoring research: when exploring inter-individual differences in fetal development, the choice of statistical method should also consider the one-versus-many approach, to avoid loss of sensitive data.

Finally, although the sample size of this study is limited, the sensitivity demonstrated by the one-versus-many framework still suggests that urinary proteomics holds promise as a low-cost, non-invasive auxiliary tool for individualized pregnancy monitoring. In future work, it is recommended to increase sampling density, expand sample size, and integrate phenotypic data such as fetal ultrasonography, which is expected to enable the establishment of a more complete temporal reference atlas of the pregnancy urinary proteome, providing molecular-level evidence for precision obstetrics and individualized fetal development assessment.

## Notes

Funding: National Key Research and Development Program of China (2023YFA1801900); Beijing Natural Science Foundation (L2604022, L246002)

### Competing Interest Statement

The authors have declared no competing interest.

## References

[1] Zheng J, Liu L, Wang J, Jin Q. Urinary proteomic and non-affinity quantitative phosphoproteomic analysis during pregnancy and non-pregnancy. BMC Genomics. 2013;14:777. DOI: 10.1186/1471-2164-14-777.

[2] Gao Y. (2013). Urine-an untapped goldmine for biomarker discovery?. Science China. Life sciences, 56(12), 1145–1146. 10.1007/s11427-013-4574-1

[3] Li L, Pan X, Wang T, Hua Y, Gao Y. Urine proteome changes in an á-synuclein transgenic mouse model of Parkinson’s disease. bioRxiv. 2020. DOI: 10.1101/2020.04.05.026104

[4] Xin, Y., Yang, Y., Qian, Q., Xia, T., Zhang, W., Dai, T., Li, R., Liu, Z., & Zhang, C. (2025). A pilot study of the plasma and urinary neurotransmitters in Chinese children with autism spectrum disorder. BMC psychiatry, 26(1), 59. 10.1186/s12888-025-07682-7

[5] Wu, J., Zhang, J., Wei, J., Zhao, Y., & Gao, Y. (2020). Urinary biomarker discovery in gliomas using mass spectrometry-based clinical proteomics. Chinese neurosurgical journal, 6, 11. 10.1186/s41016-020-00190-5

[6] Zhao, Y., Li, Y., Liu, W., Xing, S., Wang, D., Chen, J., Sun, L., Mu, J., Liu, W., Xing, B., Sun, W., & He, F. (2020). Identification of noninvasive diagnostic biomarkers for hepatocellular carcinoma by urinary proteomics. Journal of proteomics, 225, 103780. 10.1016/j.jprot.2020.103780

[7] Qin, W., Li, L., Wang, T., Huang, H., & Gao, Y. (2019). Urine Proteome Changes in a TNBS-Induced Colitis Rat Model. Proteomics. Clinical applications, 13(5), e1800100. 10.1002/prca.201800100

[8] Zhao, M., Wu, J., Li, X., & Gao, Y. (2018). Urinary candidate biomarkers in an experimental autoimmune myocarditis rat model. Journal of proteomics, 179, 71–79. 10.1016/j.jprot.2018.02.032

[9] Soma-Pillay, P., Nelson-Piercy, C., Tolppanen, H., & Mebazaa, A. (2016). Physiological changes in pregnancy. Cardiovascular journal of Africa, 27(2), 89–94. 10.5830/CVJA-2016-021

[10] Wang, X., Zhao, M., Guo, Z., Song, S., Liu, S., Yuan, T., Fu, Y., Dong, Y., Sun, H., Liu, X., Zhou, D., Zhao, W., & Sun, W. (2022). Urinary proteomic analysis during pregnancy and its potential application in early prediction of gestational diabetes mellitus and spontaneous abortion. Annals of translational medicine, 10(13), 736. 10.21037/atm-21-3497

[11] Tang, S., & Gao, Y. (2022). Urinary Proteome Changes during Pregnancy in Rats. Biomolecules, 13(1), 34. 10.3390/biom13010034

[12] Wang, H., Ge, L., Chen, S., Sun, L., Sun, W., & Gao, Y. (2025). Proteome Analysis of Daily Urine Samples of Pregnant Rats Unveils Developmental Processes of Fetus as Well as Physiological Changes in Mother Rats. Biology, 14(12), 1700. 10.3390/biology14121700

[13] Bao Y, Gao Y. Changes in the urinary proteome before and after massage in healthy individuals. ChinaXiv:202302.00108v2. 2023.

[14] Shao, C., Zhao, M., Chen, X., Sun, H., Yang, Y., Xiao, X., Guo, Z., Liu, X., Lv, Y., Chen, X., Sun, W., Wu, D., & Gao, Y. (2019). Comprehensive Analysis of Individual Variation in the Urinary Proteome Revealed Significant Gender Differences. Molecular & cellular proteomics: MCP, 18(6), 1110–1122. 10.1074/mcp.RA119.001343

[15] Sato, N., & Miyasaka, N. (2019). Heterogeneity in fetal growth velocity. Scientific reports, 9(1), 11304. 10.1038/s41598-019-47839-5

[16] Chen, R., Mias, G. I., Li-Pook-Than, J., Jiang, L., Lam, H. Y., Chen, R., Miriami, E., Karczewski, K. J., Hariharan, M., Dewey, F. E., Cheng, Y., Clark, M. J., Im, H., Habegger, L., Balasubramanian, S., O’Huallachain, M., Dudley, J. T., Hillenmeyer, S., Haraksingh, R., Sharon, D.,…Snyder, M. (2012). Personal omics profiling reveals dynamic molecular and medical phenotypes. Cell, 148(6), 1293–1307. 10.1016/j.cell.2012.02.009

[17] Li, Q., Schissler, A. G., Gardeux, V., Achour, I., Kenost, C., Berghout, J., Li, H., Zhang, H. H., & Lussier, Y. A. (2017). N-of-1-pathways MixEnrich: advancing precision medicine via single-subject analysis in discovering dynamic changes of transcriptomes. BMC medical genomics, 10(Suppl 1), 27. 10.1186/s12920-017-0263-4

